# Calcineurin promotes adaptation to chronic stress through two distinct mechanisms

**DOI:** 10.1101/2024.03.19.585797

**Authors:** Mackenzie J. Flynn, Nicholas W. Harper, Rui Li, Lihua Julie Zhu, Michael J. Lee, Jennifer A. Benanti

## Abstract

Adaptation to environmental stress requires coordination between stress-defense programs and cell cycle progression. The immediate response to many stressors has been well characterized, but how cells survive in challenging environments long-term is unknown. Here, we investigate the role of the stress-activated phosphatase calcineurin (CN) in adaptation to chronic CaCl_2_ stress in *Saccharomyces cerevisiae.* We find that prolonged exposure to CaCl_2_ impairs mitochondrial function and demonstrate that cells respond to this stressor using two CN-dependent mechanisms – one that requires the downstream transcription factor Crz1 and another that is Crz1-independent. Our data indicate that CN maintains cellular fitness by promoting cell cycle progression and preventing CaCl_2_-induced cell death. When Crz1 is present, transient CN activation suppresses cell death and promotes adaptation despite high levels of mitochondrial loss. However, in the absence of Crz1, prolonged activation of CN prevents mitochondrial loss and further cell death by upregulating glutathione (GSH) biosynthesis genes thereby mitigating damage from reactive oxygen species. These findings illustrate how cells maintain long-term fitness during chronic stress and suggest that CN promotes adaptation in challenging environments by multiple mechanisms.

## INTRODUCTION

Cells must balance the conflicting demands of proliferation and stress-defense to survive in a constantly changing environment. Under optimal growth conditions, cellular energy is predominately used to support growth and division (López-Maury *et al*., 2008; Ho and Gasch, 2015). While prioritizing proliferation ensures competitive fitness in standard conditions, rapid growth results in a reduced ability to respond to stress, which decreases fitness in sub-optimal environments (López-Maury *et al*., 2008; Ho and Gasch, 2015). Thus, in response to environmental changes, cells distribute cellular resources between growth-related processes and those providing tolerance to stress.

Slow growing cells are more resistant to stress and exhibit increased survival in challenging environments (Elliott and Futcher, 1993; Lu *et al*., 2008; Zakrzewska *et al*., 2011). Cells often induce a transient cell cycle arrest in response to stress, which provides protection against subsequent stress exposure (Bonny *et al*., 2021). While cell division is paused, cells redistribute intracellular resources to orchestrate changes in transcription, translation, and post-translational modifications (López-Maury *et al*., 2008; Ho and Gasch, 2015). In yeast, the immediate response to many stressors includes the induction of a transcriptional program called the environmental stress response (ESR) (Gasch *et al*., 2000; Causton *et al*., 2001). Approximately 300 genes required for stress-tolerance are induced by the ESR, including genes related to detoxification, cell wall integrity, and DNA damage repair. Nearly 600 genes involved in growth-related processes such as ribosome biogenesis, RNA metabolism, and nucleotide biosynthesis are also repressed by the ESR (Gasch *et al*., 2000; Causton *et al*., 2001). In addition to the ESR, which provides general protection in diverse environments, most stressors also require stress-specific changes in transcription, translation, and post-translational modifications (Gasch *et al*., 2000; Causton *et al*., 2001). As cells adapt, many of these gene expression changes are reversed and cells resume cycling. Importantly, while the immediate response to stress is well characterized, very little is known about the physiological changes that occur after cells recover from a transient stress-induced cell cycle arrest and shut off the ESR transcriptional program.

The Ca^2+^/calmodulin-activated phosphatase calcineurin (CN) is a conserved stress-response regulator. In mammals, CN is activated following the release of lysosomal Ca^2+^ and promotes lysosome biogenesis and autophagy in response to nutrient deprivation and oxidative stress (Medina *et al*., 2015; Zhang *et al*., 2016). In yeast, CN is similarly activated by the increase in cytosolic Ca^2+^ that occurs in response to many environmental stressors including cell wall damage, alkaline pH, osmotic imbalance, and toxic cations (Cyert, 2003; Cyert and Philpott, 2013). Upon activation, CN dephosphorylates target substrates involved in various cellular processes including protein trafficking, membrane structure/function, transcription, translation, cell cycle, ubiquitin signaling, and polarized growth (Goldman *et al*., 2014). The transcription factor Crz1 is the best characterized target of CN and is thought to be the major effector of the CN-dependent stress response (Stathopoulos and Cyert, 1997; Cyert, 2003). Following dephosphorylation by CN, Crz1 translocates to the nucleus and induces the expression of approximately 160 target genes with broad functions in adaptation including ion homeostasis, vesicular transport, and cell wall maintenance (Yoshimoto *et al*., 2002). Crz1 target genes also include negative regulators of CN signaling, which shut off CN activity once cells have adapted (Yoshimoto *et al*., 2002). In addition to regulating the expression of genes important for adaptation, CN delays cell cycle progression in collaboration with the MAPKs Hog1 and Mpk1 (Mizunuma *et al*., 1998, 2001; Yokoyama *et al*., 2006; Leech *et al*., 2020; Flynn and Benanti, 2022).

Many CN-dependent functions have been characterized in response to CaCl_2_ since CN is strongly activated by this stressor. However, inactivation of CN has been reported to have no effect on proliferation in the presence of moderate levels of CaCl_2_, and to confer resistance at very high concentrations (Nakamura *et al*., 1996; Matheos *et al*., 1997; Withee *et al*., 1997, 1998; Mulet *et al*., 2006; Ferreira *et al*., 2012; Xu *et al*., 2019). In contrast, Crz1 has been shown to promote proliferation in response to CaCl_2_, though reported phenotypes differ in severity (Polizotto and Cyert, 2001; Mulet *et al*., 2006; Ferreira *et al*., 2012; Zhao *et al*., 2013; Xu *et al*., 2019). Given these observations, it is unclear whether CN signaling is important for survival during CaCl_2_ stress.

Here, we characterized the physiological response of *Saccharomyces cerevisiae* to long-term growth in CaCl_2_ and investigated the impact of CN and Crz1 on cellular fitness in this environment. We show that chronic CaCl_2_ exposure triggers a decrease in proliferation rate as well as the loss of functional mitochondria. Using co-culture competitive fitness assays, we determined that CN is required for fitness during prolonged CaCl_2_ exposure and maintains fitness by both Crz1-dependent and Crz1-independent mechanisms. Our findings demonstrate how cells adapt to prolonged CaCl_2_ exposure and suggest that CN promotes cellular fitness by multiple mechanisms.

## RESULTS

### Calcineurin promotes fitness in response to chronic calcium stress

We set out to determine the importance of CN signaling for survival in CaCl_2_ stress. Consistent with previous studies, deletion of the CN-regulatory subunit *CNB1* had no detectable effect on the proliferation of cells growing on agar plates containing 0.2 M CaCl_2_ (Figure 1A) (Nakamura *et al*., 1996; Withee *et al*., 1997, 1998; Mulet *et al*., 2006; Ferreira *et al*., 2012; Xu *et al*., 2019). Since growth on plates does not always reveal modest differences in proliferation rate, we also examined the competitive growth of CN mutant strains grown in co-culture with a wild-type strain in a more sensitive assay (Conti *et al*., 2022). Briefly, one strain expressing GFP is co-cultured with a strain expressing a mutated, non-fluorescent GFP(Y66F) (Supplemental Figure S1A). This co-culture is maintained in logarithmic phase by dilution at regular intervals and the opposing GFP markers allow the relative percentages of each strain to be monitored over time by flow cytometry. Importantly, we observe equivalent fitness in strains expressing both wild-type GFP and non-fluorescent GFP(Y66F) (Supplemental Figure S1, B and C). Using this approach, we found that deletion of *CNB1* resulted in a modest proliferative defect under optimal growth conditions consistent with CN having few defined functions in the absence of stress (Figure 1B) (Cyert, 2003; Cyert and Philpott, 2013). In contrast, there was a much stronger fitness defect in *cnb11* mutant cells when co-cultured with wild-type in the presence of CaCl_2_, indicating that CN is required for fitness in this condition (Figure 1C).

**Figure 1.**
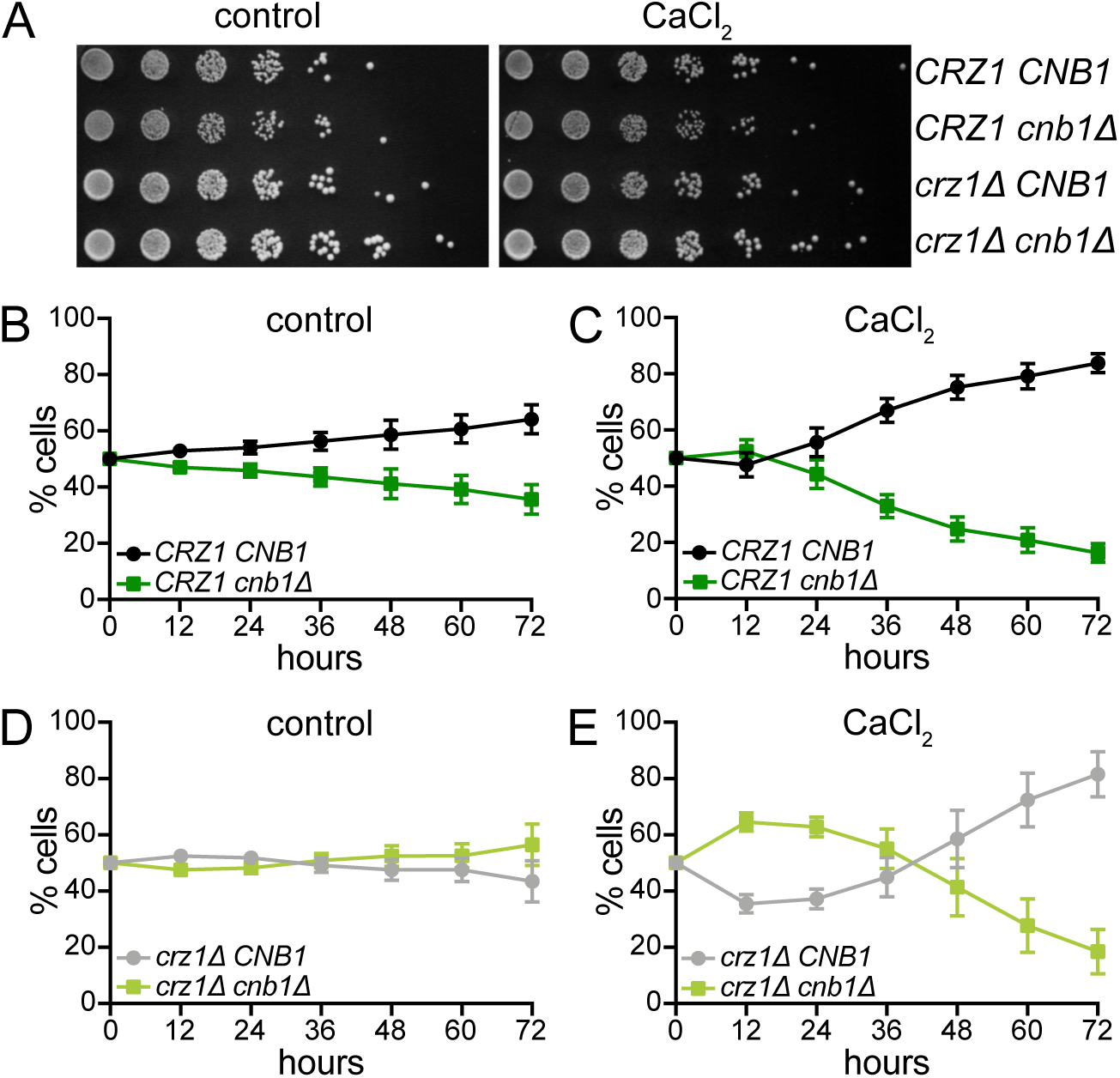
Calcineurin promotes fitness in response to chronic calcium stress. (A) Five-fold serial dilutions of the indicated strains were spotted on synthetic complete media with or without 0.2 M CaCl_2_. Plates were imaged after 72 hours of growth at 30°C. (B-C) *CRZ1 CNB1* and *CRZ1 cnb1Δ* cells were co-cultured in the absence (B) or presence (C) of 0.2 M CaCl_2_. Co-cultures were sampled and diluted every 12 hours and the percentage of each strain was determined at the indicated timepoints by flow cytometry. An average of n = 11 biological replicates is shown. Error bars indicate standard deviations. (D-E) Same as (B-C) in *crz1Δ CNB1* and *crz1Δ cnb1Δ* cells. An average of n = 9 biological replicates is shown. Error bars indicate standard deviations.

The transcription factor Crz1 is activated by CN and regulates the expression of genes important for adaptation to CaCl_2_ (Stathopoulos-Gerontides *et al*., 1999; Yoshimoto *et al*., 2002). Notably, Crz1 is known to impact fitness in response to CaCl_2_ though the severity of Crz1-dependent growth phenotypes differs between studies (Figure 1A and Supplemental Figure S1, D and E) (Polizotto and Cyert, 2001; Mulet *et al*., 2006; Ferreira *et al*., 2012; Zhao *et al*., 2013; Xu *et al*., 2019; Hsu *et al*., 2021). This raised the possibility that the CN-dependent fitness defect that we observed resulted from a failure to activate Crz1. If this is true, then *crz1Δ* and *crz11 cnb11* cells should have equivalent fitness in the presence and absence of CaCl_2_. Interestingly, while the *crz1Δ* and *crz11 cnb11* mutants had equal fitness under optimal growth conditions, CN-dependent fitness differences were observed when these strains were co-cultured in the presence of CaCl_2_ (Figure 1, D and E). Notably, *cnb1Δ* and *crz11 cnb11* cells did not display fitness defects within the first 12 hours of CaCl_2_ treatment (Figure 1, C and E), which is consistent with fact that CN promotes a transient cell cycle arrest immediately following exposure to CaCl_2_ (Leech *et al*., 2020). This arrest is lengthened in the absence of *CRZ1* and, accordingly, the *crz11 cnb11* mutant exhibited a more pronounced growth advantage during the first 12 hours of CaCl_2_ exposure (Figure 1E). Despite these early fitness benefits, both *cnb1Δ* and *crz11 cnb11* cells were depleted from the co-cultures after 12 hours indicating that CN maintains fitness during chronic CaCl_2_ in both the presence and absence of Crz1 (Figure 1, C and E). However, the Crz1-dependent and Crz1-independent mechanisms by which CN promotes fitness are unknown.

### Crz1 is required to inactivate CN

CN is strongly activated within minutes of exposure to CaCl_2_ and is subsequently inactivated as cells adapt (Stathopoulos-Gerontides *et al*., 1999; Leech *et al*., 2020; Flynn and Benanti, 2022). However, the timing of CN inactivation and adaptation during prolonged stress exposure is unclear. To better understand the response to chronic stress, we monitored CN activity as cells grew in CaCl_2_ for 72 hours using a fluorescent reporter (CNR-C), which measures CN-dependent dephosphorylation and nuclear localization of a GFP-tagged, non-functional fragment of Crz1. Consistent with previous studies (Stathopoulos-Gerontides *et al*., 1999; Leech *et al*., 2020; Flynn and Benanti, 2022), we observed CN-dependent changes in CNR-C within 10 minutes of CaCl_2_ treatment and both wild-type and *crz1Δ* cells initially activated CN to similar extents, as shown by the rapid appearance of the higher mobility CNR-C species and increased nuclear signal (Figure 2, A and B). In wild-type cells, CN was inactivated within 3 hours after CaCl_2_ treatment as indicated by the reappearance of the phosphorylated, lower mobility CNR-C species and loss of nuclear signal (Figure 2, A and B). However, even 72 hours after CaCl_2_ exposure, CN was still active in 40-50% of *crz1Δ* cells (Figure 2, C and D), consistent with Crz1 acting in a negative feedback loop to shut off CN signaling (Cyert, 2003). This observation demonstrates that Crz1 is required to inactivate CN and suggests that cells are unable to properly adapt to CaCl_2_ in the absence of Crz1.

**Figure 2.**
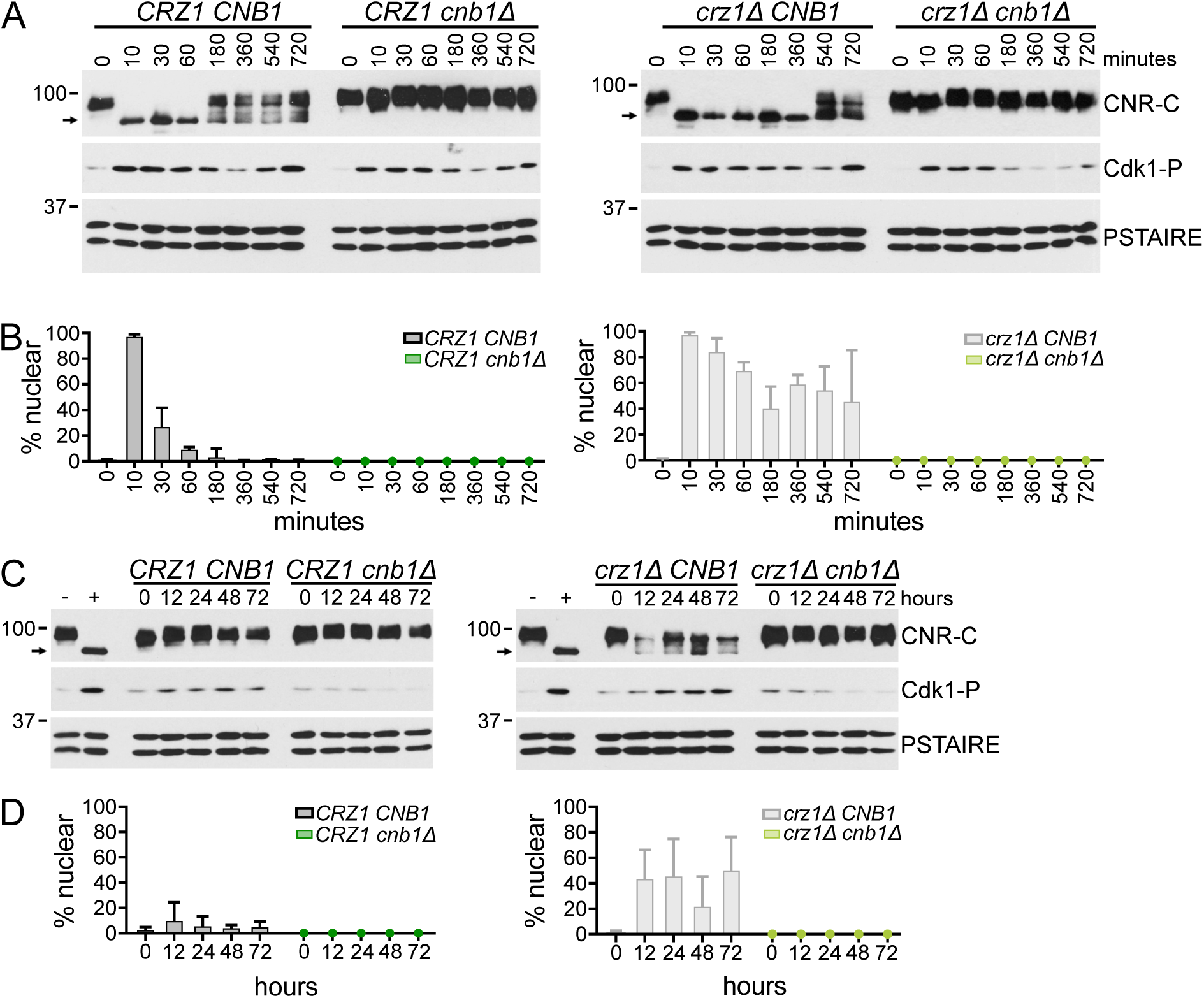
Crz1 is required to inactivate CN. (A) *CRZ1 CNB1, CRZ1 cnb1Δ*, *crz1Δ CNB1,* and *crz1Δ cnb1Δ* cells were grown in media with or without 0.2 M CaCl_2_. Samples were collected at the indicated times and Western blots were performed for CNR-C (Crz1^14-424^-GFP), Cdk1 phosphorylated on Y19 (Cdk1-P), and PSTAIRE (loading control). Arrows indicate dephosphorylated CNR-C. (B) Cells from (A) were prepared for imaging as described in the Materials and Methods and the percentage of cells with nuclear CNR-C signal was calculated. An average of n = 3 biological replicates is shown, error bars indicate standard deviations. *CRZ1 CNB1, CRZ1 cnb1Δ*, *crz1Δ CNB1,* and *crz1Δ cnb1Δ* cells were grown in media with or without 0.2 M CaCl_2_ and diluted every 12 hours. Samples were collected at the indicated times and Western blots were performed as in (A). Arrows indicate dephosphorylated CNR-C. As a control for CNR-C dephosphorylation, wild-type cells before, “-”, and after, “+”,10 minutes of 0.2 M CaCl_2_ treatment are shown. Cells from (C) were prepared as in (B). An average of n = 9 biological replicates in shown, error bars indicate standard deviations.

In response to acute CaCl_2_ stress, CN delays progression through multiple phases of the cell cycle in combination with the stress-activated MAPKs Hog1 and Mpk1 (Mizunuma *et al*., 1998, 2001; Yokoyama *et al*., 2006; Leech *et al*., 2020; Flynn and Benanti, 2022). This suggested that the fitness defects of CN mutant cells (Figure 1, C and E) may be due to slowed cell cycle progression. One way that CN delays the cell cycle is by inhibiting Cdk1 activity. Activation of CN, alongside Hog1 and Mpk1, indirectly activates the kinase Swe1, which then phosphorylates Cdk1 on Y19 (Cdk1-P) to inhibit Cdk1 activity and delay cell cycle progression (Mizunuma *et al*., 1998, 2001; Clotet *et al*., 2006). As previously reported (Mizunuma *et al*., 1998, 2001; Leech *et al*., 2020), exposure to CaCl_2_ triggered an increase in Cdk1-P and a transient cell cycle arrest (Figure 2A and Supplemental Figure S2). Immediately following CaCl_2_ treatment, levels of Cdk1-P were comparable in all genotypes consistent with redundant control of Cdk1 inhibition during stress (Mizunuma *et al*., 1998, 2001; Clotet *et al*., 2006). However, CN was required to maintain the inhibitory phosphorylation on Cdk1 at time points longer than 12 hours (Figure 2C). Since cells lacking Cnb1 display a proliferative defect in competitive growth assays (Figure 1, C and E), this suggests that the observed proliferation defect is not the result of increased Cdk1 inhibition. Consequently, CN must regulate additional processes to promote fitness during chronic stress.

### Gene expression changes in response to chronic calcium stress

To understand the physiological changes that cells undergo in response to chronic stress, we used RNA-seq to measure gene expression during long-term growth in CaCl_2_ (Supplemental Figure S3A). To determine the role of CN and Crz1 in this process, we also examined gene expression in the *cnb11, crz1Δ,* and *crz11 cnb11* mutants over 72 hours of CaCl_2_ treatment. In response to CaCl_2_, approximately half of the genome was differentially expressed at one or more timepoints and these genes were highly overlapping across genotypes (Supplemental Figure S3, B and C). This was in stark contrast to the changes that occurred in response to chronic KCl treatment, a stressor that does not require CN for fitness, in which fewer than 200 genes were differentially expressed (Supplemental Figure S4, A-C, Supplemental Data File S1). To examine the categories of genes changing in expression over time in response to CaCl_2_, we applied hierarchical clustering and found that, in all genotypes, differentially expressed genes fell into two broad clusters of genes whose expression either increased or decreased over time (Supplemental Figure S3D, Supplemental Data File S2). Gene Ontology (GO)-term enrichment analysis of upregulated and downregulated clusters in each genotype revealed several commonly regulated biological processes (Supplemental Figure S4, E-H, Supplemental Data File S2). Similar results were obtained using Gene Set Enrichment Analysis (GSEA) to identify gene sets that were significantly correlated with each the genotype (Figure 3, Supplemental Data File S3). The most notable processes that changed were related to translation/ribosome biogenesis, amino acid synthesis, ion transport, and mitochondrial function (Figure 3 and Supplemental Figure S3, E-H).

**Figure 3.**
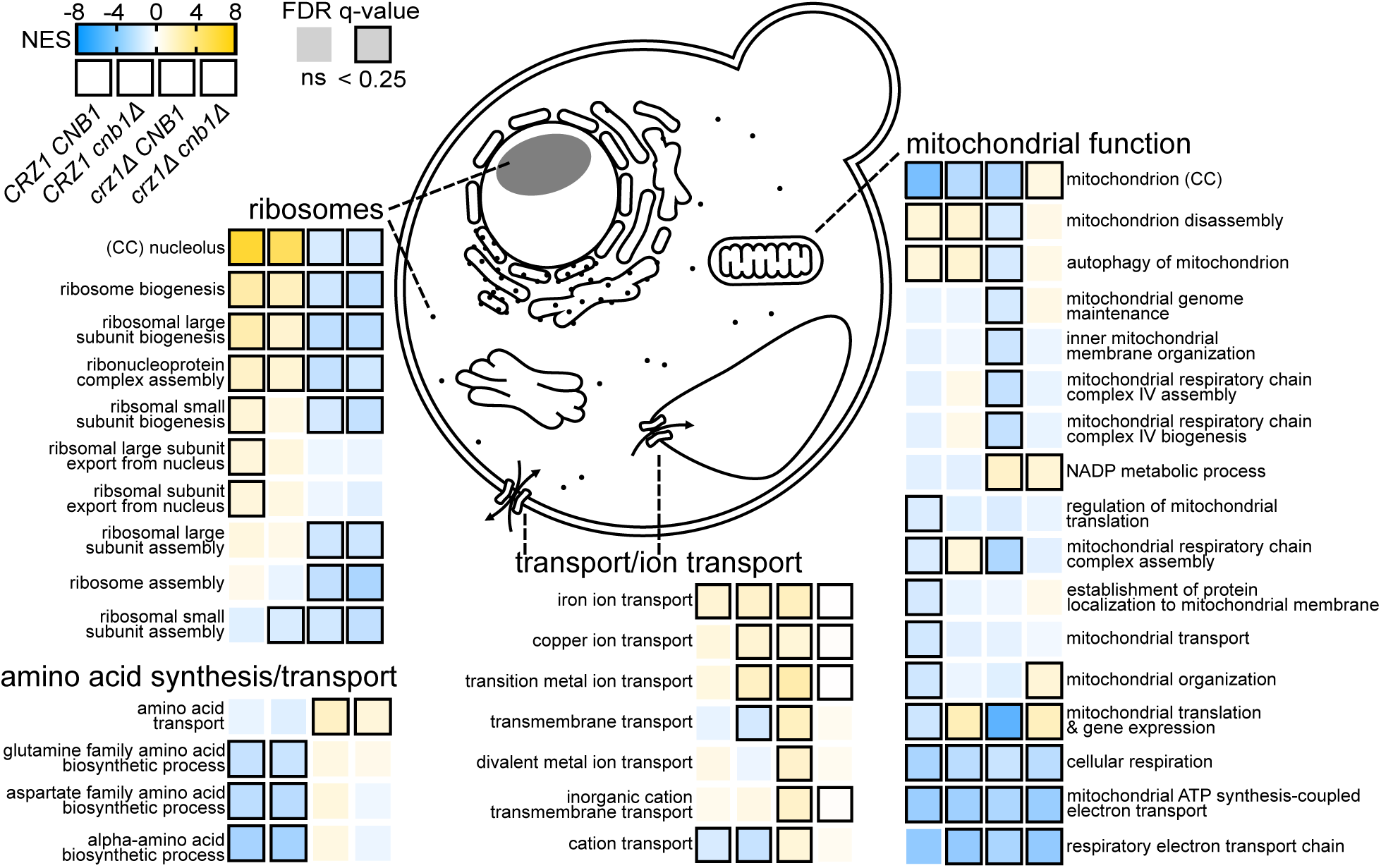
Gene expression changes in response to chronic calcium stress. Gene Set Enrichment Analysis (GSEA) of differentially expressed genes from *CRZ1 CNB1, CRZ1 cnb1Δ*, *crz1Δ CNB1,* and *crz1Δ cnb1Δ* strains after 72 hours of growth in 0.2 M CaCl_2_, compared to genotype-matched unstressed controls. All categories are ‘GO Biological Process’ with the exception of those marked ‘CC’ for ‘GO Cellular Component’. Each square represents a genotype and colors indicate the normalized enrichment score (NES). Outlined squares indicate significantly enriched (FDR q-value < 0.25) categories. GSEA data for all genotypes at all timepoints is included in Supplemental Data File S3.

One striking feature of the response to chronic CaCl_2_ is how it differed from the well-defined ESR transcriptional program (Supplemental Figure S4D) (Gasch *et al*., 2000; Causton *et al*., 2001). For example, genes involved in translation and ribosome biogenesis, which are downregulated in the ESR immediately following stress (Gasch *et al*., 2000; Causton *et al*., 2001), were upregulated during chronic CaCl_2_ stress (Figure 3 and Supplemental Figure S3, E-H). Interestingly, upregulation of ribosome biogenesis genes was dependent upon Crz1 but not CN, since these categories of genes were depleted in both *CNB1* and *cnb11* cells lacking Crz1. Conversely, genes involved in processes related to amino acid biosynthesis and transport were significantly downregulated in a Crz1-dependent manner.

In addition to categories displaying Crz1-dependent enrichment or depletion, some processes shared common regulation across all genotypes (Figure 3 and Supplemental Figure S3, D-H). For example, we observed enrichment among upregulated genes for processes related to ion transport. Many of these genes are involved in iron transport across the plasma, vacuolar, and endoplasmic reticulum membranes, indicating that regulation of iron availability may be important during adaptation to chronic CaCl_2_ stress. Lastly, processes related to mitochondrial function including mitochondrial translation, cellular respiration, and the electron transport chain were enriched among downregulated genes in all genotypes (Figure 3 and Supplemental Figure S3, E-H). This finding highlights another way in which chronic stress differs from the ESR, since mitochondrial function is upregulated in the ESR during acute stress exposure (Gasch *et al*., 2000; Causton *et al*., 2001).

### CN and Crz1 regulate translation, mitochondrial loss, and viability during prolonged stress

Our data demonstrate that ribosome biogenesis genes are upregulated in *CRZ1* cells independent of CN (Figure 4A). We hypothesized that *CRZ1* cells may have increased translational capacity following exposure to CaCl_2_ and should therefore be more resistant to translational inhibition. To test this, we evaluated growth in the presence of sub-lethal doses of the translation inhibitor cycloheximide (CHX). Following 72 hours of growth in CaCl_2_, *CRZ1* cells were more tolerant to CHX compared to cells lacking Crz1 as well as identical strains that were not treated with CaCl_2_ (Figure 4B). These data indicate that Crz1 increases the translational capacity of cells in response to CaCl_2_ through a CN-independent mechanism and suggest that cells may require more ribosomes to survive and proliferate in chronic stress.

**Figure 4.**
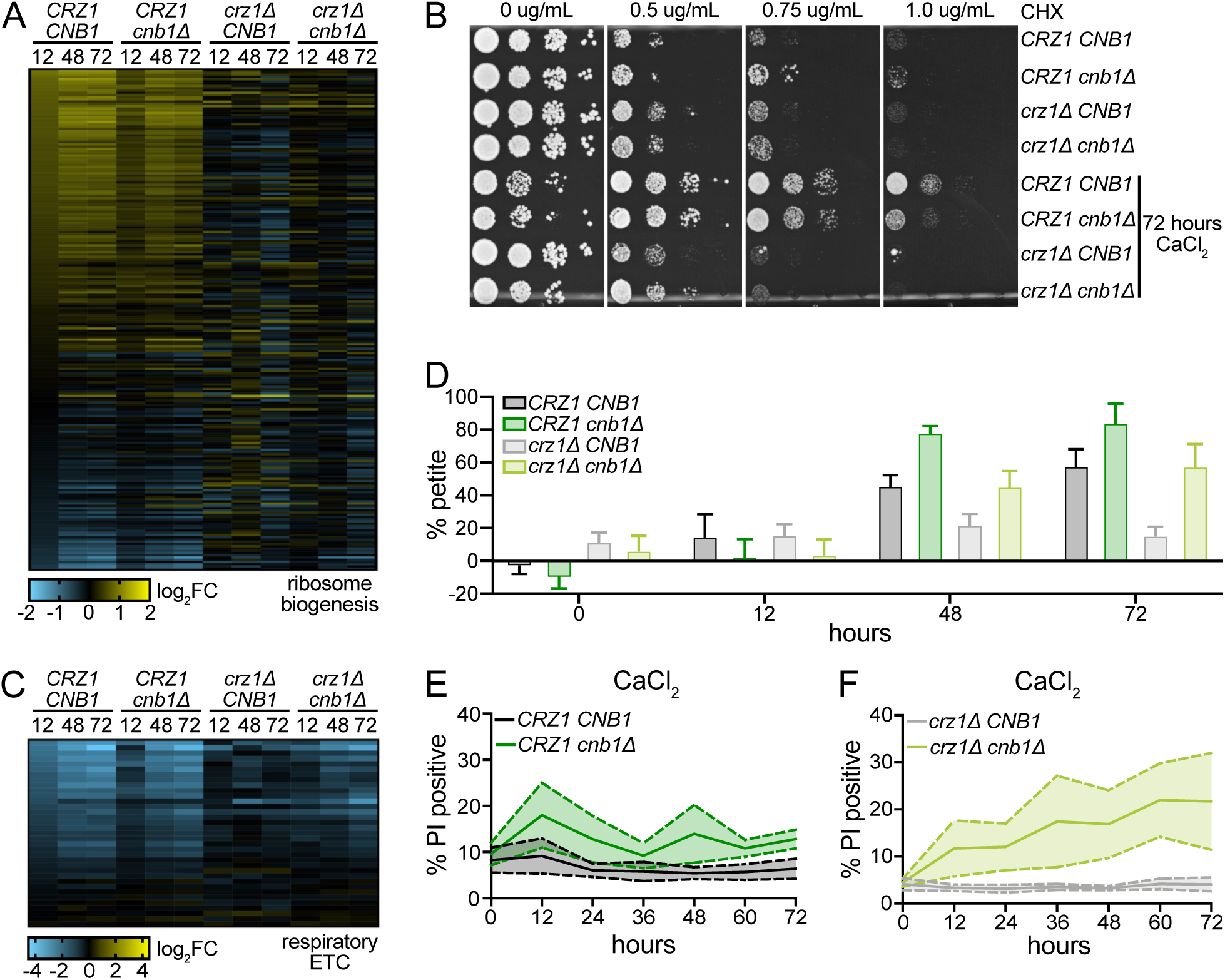
CN and Crz1 regulate translation, mitochondrial loss, and viability during prolonged stress. (A) Heat map showing the log_2_ fold change values for all genes included in ‘ribosome biogenesis’ (GO:0042254) at the indicated timepoints compared to an unstressed control for each designated genotype. A list of genes and log_2_ fold change values used to generate the heatmap are included in Supplemental Data File S4. (B) After 72 hours of growth in 0.2 M CaCl_2_, ten-fold serial dilutions of the indicated strains were spotted onto YPD agar containing the specified concentrations of cycloheximide (CHX). Unstressed controls for each strain are included for comparison. Representative images from n = 3 experiments are shown. (C) As in (A) for all genes included in ‘respiratory electron transport chain’ (GO:0022904). A list of genes and log_2_ fold change values used to generate the heatmap are included in Supplemental Data File S4. (D) *CRZ1 CNB1*, *CRZ1 cnb1Δ, crz1Δ CNB1,* and *crz1Δ cnb1Δ* cells were grown for the indicated times in 0.2 M CaCl_2_ prior to measuring petite frequency as described in the Materials and Methods. An average of n = 6 biological replicates is shown. Error bars indicate standard deviations. (E-F) *CRZ1 CNB1* and *CRZ1 cnb1Δ* (E) or *crz1Δ CNB1* and *crz1Δ cnb1Δ* (F) cells were grown in the presence of 0.2 M CaCl_2_. Cultures were sampled and diluted at the indicated times and the proportion of propidium iodide (PI) positive cells was determined by flow cytometry. Shown is an average of n = 12 (E) or n = 9 (F) biological replicates. Solid line denotes the mean and the shaded area within the dashed lines denotes the standard error of the mean.

Our gene expression data also predict that mitochondrial function is reduced during chronic CaCl_2_ exposure, since the expression of genes encoding proteins with mitochondrial functions is decreased in all genotypes (Figure 3 and Supplemental Figure S3, E-H). A critical role of the mitochondria is to produce ATP via oxidative phosphorylation (Alberts *et al*., 1994). In agreement with the GSEA and GO-term analyses, we observed CaCl_2_-dependent downregulation of genes involved in the electron transport chain (ETC) (Figure 4C). While ETC transcripts decreased in all genotypes following the addition of CaCl_2_, this downregulation was less severe in cells lacking Crz1. Decreased expression of ETC genes has been previously linked to the loss of functional mitochondria (Epstein *et al*., 2001). Consistent with that study, we observed an increased proportion of petite colonies that cannot grow on non-fermentable carbon sources in all genotypes except the *crz1Δ* mutant, which maintains active CN throughout the time course (Figure 4D, Figure 2). Notably, changes in petite frequency generally correlated with the degree of ETC transcript downregulation in each genotype (Figure 4C). These data indicate that chronic CaCl_2_ stress triggers mitochondrial dysfunction and suggest that CN activity helps prevent mitochondrial loss.

Increases in cytosolic calcium concentrations have been shown to precede mitochondrial permeabilization, which impairs mitochondrial function and triggers cell death in both yeast and mammals (Guaragnella *et al*., 2012; Carraro and Bernardi, 2016; Frigo *et al*., 2023). Since chronic growth in CaCl_2_ compromises mitochondrial function (Figures 2 and 4, C and D), we next monitored cell viability using propidium iodide to determine if CaCl_2_ triggered corresponding increases in cell death (Figure 4E) (Deere *et al*., 1998; Mirisola *et al*., 2014; Carmona-Gutierrez *et al*., 2018). Although CN did not impact viability in the absence of stress (Supplemental Figure S5A), we observed a CN-dependent increase in cell death after 12 hours in CaCl_2_ that steadily decreased to near pre-stress levels by the 72-hour timepoint (Figure 4E). This finding suggests that CN normally suppresses CaCl_2_-induced death, however additional mechanisms may preserve viability at later timepoints.

To examine the role of Crz1 in this process, we measured cell death in *crz1Δ* and *crz11 cnb11* cells grown in the presence of CaCl_2_. Similar to what we observed in the *cnb11* single mutant (Figure 4F and Supplemental Figure S5B), the proportion of PI-positive cells increased in the *crz11 cnb11* double mutant after 12 hours in CaCl_2_, demonstrating that CN inhibits CaCl_2_-induced death independent of Crz1 at early timepoints. However, death in the double mutant continued to increase for the remainder of the experiment (Figure 4F). This suggests that cells may be unable to adapt in the absence of Crz1, and prolonged CN activity in cells lacking Crz1 suppresses further CaCl_2_-induced death (Figure 2). Taken together these data suggest that both CN and Crz1 help maintain cell viability during chronic stress exposure.

### CN stimulates prolonged expression of glutathione biosynthesis genes in the absence of Crz1

Prolonged exposure to CaCl_2_ stress impairs mitochondrial function and triggers CN-dependent cell death (Figure 4, C-F). Mitochondrial dysfunction, as well as exposure to various environmental stressors, has been shown to increase intracellular reactive oxygen species (ROS) (Brennan and Schiestl, 1996; Davidson *et al*., 1996; Granot *et al*., 2003; Drakulic *et al*., 2005; Du *et al*., 2007; Dudgeon *et al*., 2008; Yi *et al*., 2018; Bothammal *et al*., 2022; Trip *et al*., 2022). To protect against ROS-induced damage to proteins, lipids, and nucleic acids, cells rely on the antioxidant glutathione (GSH) (Jamieson, 1998). Notably, activation of CN promotes the expression of several genes required for GSH biosynthesis following acute exposure to CaCl_2_ stress (Yoshimoto *et al*., 2002), suggesting that CN may increase GSH production to prevent mitochondrial loss and cell death.

To determine if GSH levels are regulated in a CN-dependent manner during chronic CaCl_2_ stress we examined the expression of genes required to synthesize GSH (Figure 5A). In wild type and *cnb11* cells, there was either no change or a modest decrease in the GSH biosynthesis genes following 12 to 72 hours of growth in CaCl_2_, consistent with CN being inactive in these strains at these timepoints (Figure 5, B and C, Figure 2, C and D). In contrast, *crz1Δ* cells, in which CN remains active, maintained elevated expression of many GSH biosynthetic genes in a CN-dependent manner (Figure 5, B and C). Interestingly, these gene expression changes are correlated with the fact that *crz1Δ* cells are protected from mitochondrial loss and downregulation of ETC genes (Figures 5, B and C, and 4, C and D). These findings suggest that CN promotes expression of GSH biosynthesis genes in response to CaCl_2_, and prolonged CN activity in the absence of Crz1 maintains this pattern of gene expression.

**Figure 5.**
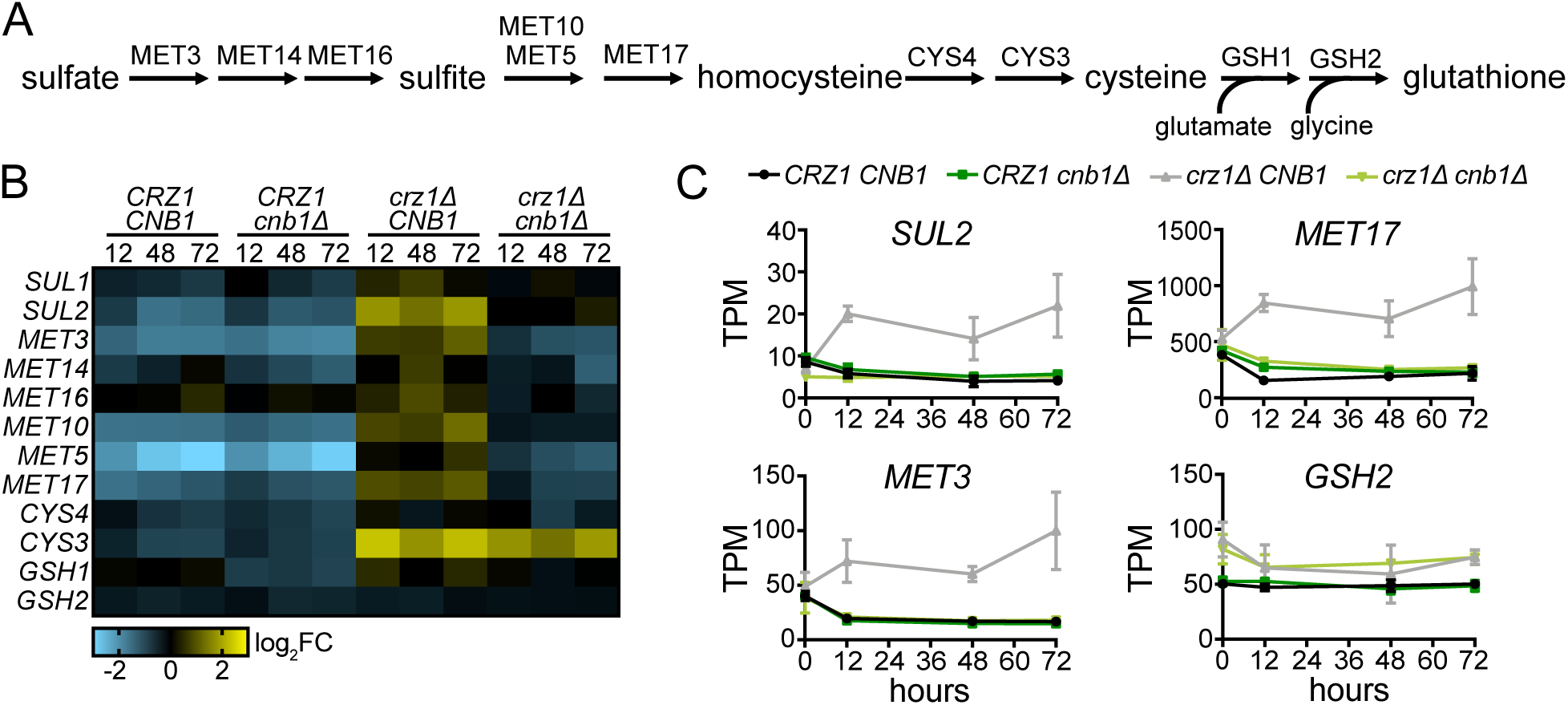
CN stimulates prolonged expression of glutathione biosynthesis genes in the absence of Crz1. (A) Schematic of glutathione biosynthesis in *S. cerevisiae.* (B) Heatmap showing the log_2_ fold change in expression of genes involved in glutathione production compared to a corresponding unstressed control in each genotype. A list of genes and log_2_ fold change values used to generate the heatmap are included in Supplemental Data File S4. (C) TPM values at the indicated timepoints for *SUL2, MET3, MET17,* and *GSH2* in *CRZ1 CNB1*, *CRZ1 cnb1Δ, crz1Δ CNB1,* and *crz1Δ cnb1Δ* cells. An average of n = 3 biological replicates is shown, error bars indicate standard deviations.

### Antioxidants rescue fitness and cell death in the absence of CN and Crz1

Having established that prolonged CN activity in the absence of Crz1 sustains the expression of GSH biosynthetic genes, we next wanted to determine if the CN-dependent upregulation of these genes protects cells from ROS-induced death. To test this hypothesis, we first examined the competitive growth of the *crz11 cnb11* strain relative to a *crz1Δ* strain, in the presence of both CaCl_2_ and GSH (Figure 6A). Similar to what we observed in response to CaCl_2_ alone, c*rz11 cnb11* cells had a fitness benefit compared to the c*rz1Δ* mutant in the first 12 hours (compare Figures 1E and 6A). However, unlike during treatment with only CaCl_2_, the c*rz11 cnb11* double mutant maintained this early proliferative advantage in the presence GSH. This result is consistent with our hypothesis that the prolonged CN activity in *crz1Δ* cells upregulates the expression of GSH biosynthetic genes to promote fitness (Figures 6A and 5, B and C). In addition to rescuing the CaCl_2_-induced fitness defect of the *crz11 cnb11* mutant, GSH also reduced cell death, further supporting a GSH-dependent rescue of fitness (Figure 6B). A similar rescue of fitness and cell death was observed with homocysteine (HCys), an upstream metabolite in the GSH synthesis pathway (Figure 5A and 6, C and D), suggesting that CN promotes fitness through a GSH-dependent mechanism when Crz1 is absent.

**Figure 6.**
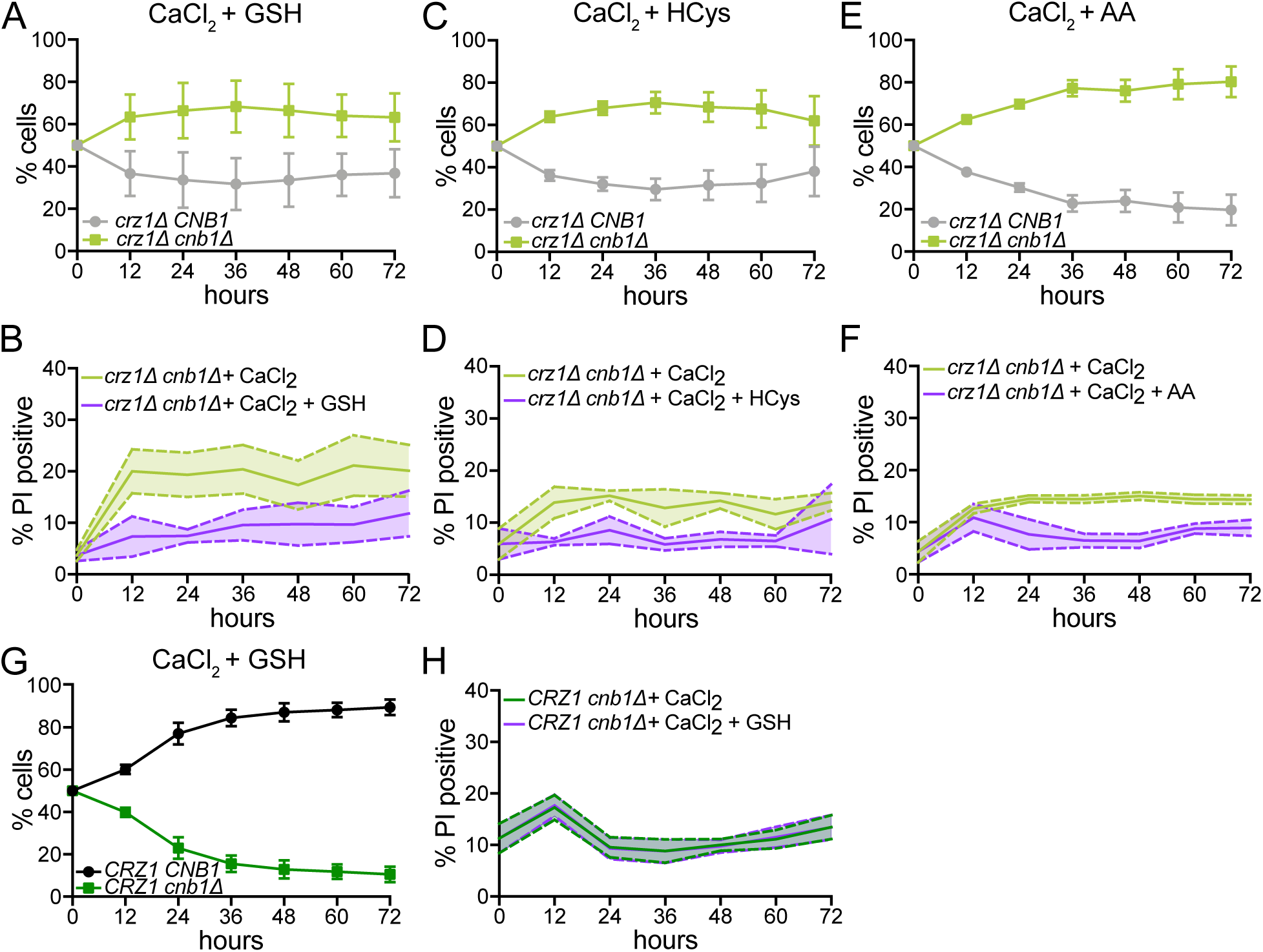
Antioxidants rescue fitness and cell death in the absence of CN and Crz1. (A) *crz1Δ CNB1* and *crz1Δ cnb1Δ* cells were co-cultured in the presence of 0.2 M CaCl_2_ and 250 µM reduced glutathione (GSH). Co-cultures were sampled and diluted every 12 hours and the percentage of each strain was quantified at the indicated times. An average of n = 9 biological replicates is shown. Error bars indicate standard deviations. (B) *crz1Δ cnb1Δ* cells were grown in the presence of 0.2 M CaCl_2_ or 0.2 M CaCl_2_ and 250 µM GSH and diluted every 12 hours. The percentage of PI-positive cells was measured by flow cytometry. An average of n = 9 biological replicates is shown. Solid line denotes the mean and the shaded area within the dashed lines denotes the standard error of the mean. (C) As in (A) in the presence of 0.2 M CaCl_2_ and 250 µM homocysteine (HCys). An average of n = 6 biological replicates is shown. Error bars indicate standard deviations. (D) As in (B) in the presence of 0.2 M CaCl_2_ and 250 µM homocysteine. An average of n = 6 biological replicates is shown. Solid line denotes the mean and the shaded area between the dashed lines denotes the standard error of the mean. (E) As in (A) in presence of 0.2 M CaCl_2_ and 5 mM ascorbic acid (AA). An average of n = 6 biological replicates is shown. Error bars indicate standard deviations. (F) As in (B) in the presence of 0.2 M CaCl_2_ and 5 mM AA. An average of n = 6 biological replicates is shown. Solid line denotes the mean and the shaded area between the dashed lines denotes the standard error of the mean. (G) Same as (A) in *CRZ1 CNB1* and *CRZ1 cnb1Δ* cells. An average of n = 9 biological replicates is shown. Error bars indicate standard deviations. (B) Same as (B) in *CRZ1 cnb1Δ* cells. An average of n = 12 biological replicates is shown. Solid line denotes the mean and the shaded area within the dashed lines denotes the standard error of the mean.

In addition to being an antioxidant, GSH is used as a source of intracellular sulfur to produce the amino acids cysteine and methionine (Elskens *et al*., 1991). Thus, supplementing cells with GSH or HCys could impact fitness and viability by either maintaining redox homeostasis or supporting amino acid production. To discriminate between these possibilities, we measured fitness and death in cells supplemented with the antioxidant ascorbic acid (AA), which is structurally distinct from GSH and cannot be used as a source of sulfur (Jamieson, 1998). Comparable to what we observe with GSH or HCys, AA rescued CaCl_2_-induced fitness defects and prevented death in the *crz11 cnb11* mutant (Figure 6, E and F). Together these data demonstrate that GSH rescues fitness due to its role as an antioxidant and suggest that *crz11 cnb11* cells may be dying as a result of oxidative damage.

We next examined the effect of GSH on fitness and viability in *CRZ1* cells. Since the expression of GSH synthesis genes was comparable between wild-type and *cnb11* cells, we did not expect the fitness of *cnb11* cells to be improved by the addition of GSH. As predicted, the *cnb11* mutant exhibited a pronounced fitness defect in the presence of both CaCl_2_ and GSH (Figure 6G), similar to what was observed in response to CaCl_2_ alone (Figure 1C). Likewise, the increase in CaCl_2_-induced cell death that occurred in *cnb11* cells after 12 hours did not change upon treatment with GSH (Figure 6H). These results indicate that CN promotes fitness and viability in a GSH-independent manner in cells expressing Crz1.

### Altered proliferation and death differentially contribute to fitness in CN mutants

Our data suggest that CN impacts both cell cycle progression and cell death during chronic CaCl_2_ stress (Figures 2, A and C, and 4, E and F, and Supplemental Figure S2, A-D). However, the relative contributions of proliferation and death to CN-dependent fitness cannot be determined by measuring net population growth since it is a combination of proliferation and cell death. To determine the contributions of each, we measured the net population growth and the percentage of dead cells in monocultures growing in the presence of CaCl_2_ (Figure 7, A-D). We then used these measurements to calculate the death rates and true proliferation rates, which excludes influences from cell death, for each strain (Schwartz *et al*., 2020). After 12 hours in CaCl_2_, we observed a decrease in proliferation rate in all genotypes. However, while the proliferation rate mostly stabilized in cells with functional CN after 12 hours, the proliferation rate of cells lacking Cnb1 continued to decrease for the remainder of the time course. We also measured a CN-dependent increase in the CaCl_2_-induced death rate (Figure 7, A-D), consistent with the results of the PI staining (Figure 4, E and F). These findings suggest that the CN-dependent fitness defects we observed in the co-culture competitive growth assays (Figure 1, C and E) are due to both a reduced rate of cell division and an increased rate of death.

**Figure 7.**
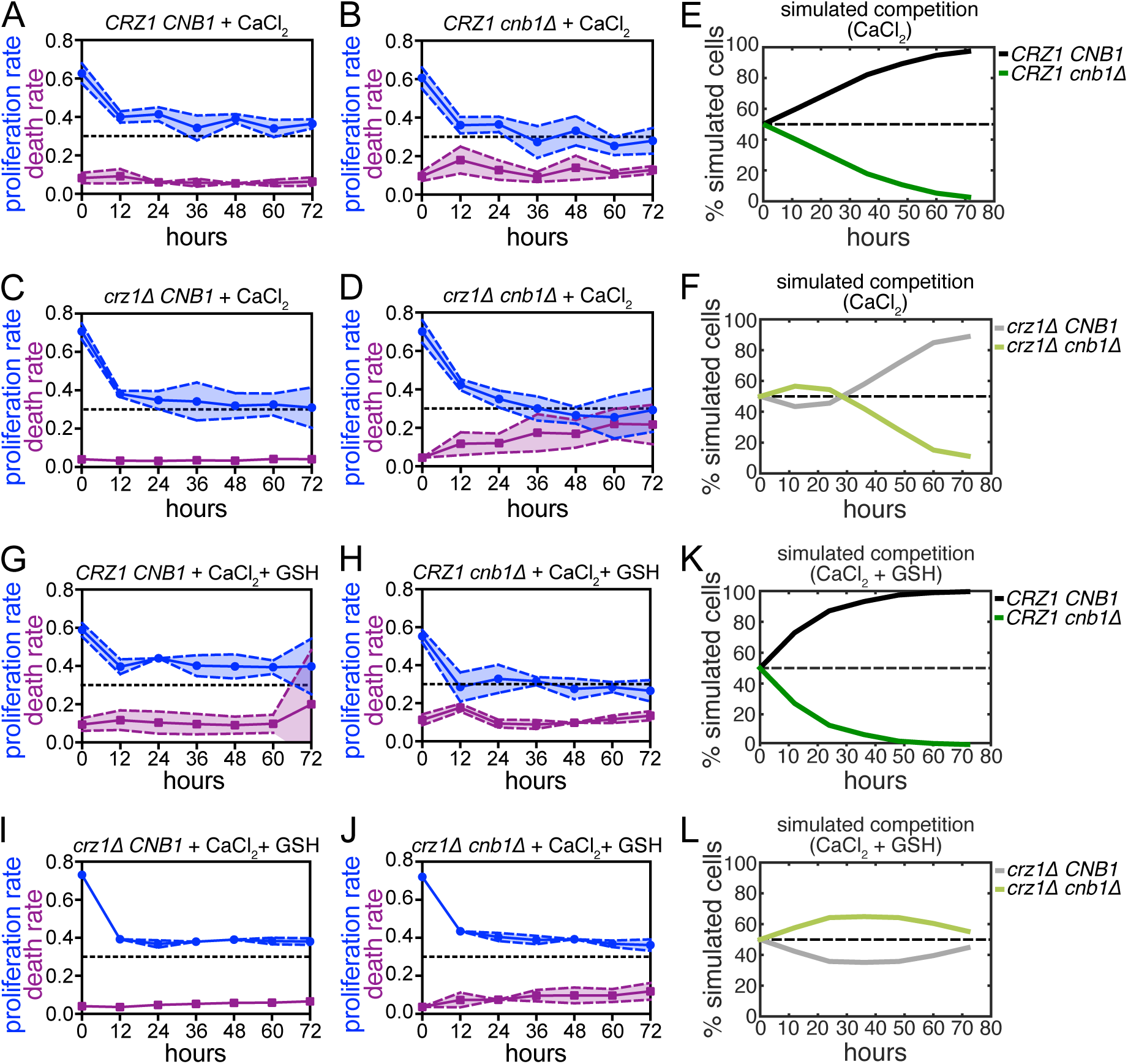
Altered proliferation and death differentially contribute to fitness in CN mutants. (A-D) Proliferation and death rates of *CRZ1 CNB1* (A), *CRZ1 cnb1Δ* (B), *crz1Δ CNB1* (C), and *crz1Δ cnb1Δ* (D) cells growing in 0.2 M CaCl_2_ calculated from data presented in Figure 4E (A-B) and Figure 4F (C-D). (E-F) Proliferation and death rates determined in (A-D) were used to simulate co-culture competition assays between *CRZ1 CNB1* and *CRZ1 cnb1Δ* (E) or *crz1Δ CNB1* and *crz1Δ cnb1Δ* (F) cells. (G-J) Proliferation and death rates of *CRZ1 CNB1* (G), *CRZ1 cnb1Δ* (H), *crz1Δ CNB1* (I), and *crz1Δ cnb1Δ* (J) cells growing in 0.2 M CaCl_2_ + 250 µM GSH calculated from data presented in Supplemental Figure S5C (G), Figure 6H (H), Supplemental Figure S5D (I), and Figure 6B (J). (K-L) Proliferation and death rates determined in (G-J) were used to simulate co-culture competition assays between *CRZ1 CNB1* and *CRZ1 cnb1Δ* (K) or *crz1Δ CNB1* and *crz1Δ cnb1Δ* (L) cells.

To better understand the relationship between proliferation and viability, we next used the proliferation and death rates determined from cells growing in monoculture to simulate the co-culture competitive growth assays (Figure 7, E and F). Notably, this simulated growth model did not consider cell non-autonomous interactions. In doing this, we were able to recapitulate the CN-dependent fitness defects that we observed in both the presence and absence of Crz1 (Figure 1). Importantly, this finding demonstrates that our estimations of proliferation and death rates from monoculture data accurately reflect the fitness dynamics of co-cultured cells. Moreover, it suggests that the effects of each genotype are cell autonomous and co-culture growth is not impacted by any secreted molecules, such as GSH.

To determine whether CN affects fitness through different mechanisms in the presence and absence of Crz1, we examined the relative contributions of decreased proliferation rate and increased cell death in each genotype. To do this, we performed additional co-culture simulations in which we fixed the proliferation or death rates of the competing strains (Supplemental Figure S6, A-D). We first examined fitness in the *cnb1Δ* mutant relative to wild-type cells. When the death rates, but not the proliferation rates, of wild-type and *cnb11* cells were fixed the model more closely replicated the experimentally determined fitness phenotype (Supplemental Figure S6, A and B). This result indicates that a proliferative defect is the primary factor decreasing fitness of the *cnb1Δ* mutant. In contrast, cell death appears to contribute more to the fitness defects in the CN mutant in the absence of Crz1 (Figure 7F and Supplemental Figure S6, C and D). When either the proliferation or death rates of the *crz1Δ* and *crz11 cnb11* mutants were fixed in simulations, partial fitness defects were observed in the double mutant (Supplemental Figure S6, C and D). This suggests that fitness, while predominantly driven by decreased proliferation, is amplified by the relatively high levels of death in the absence of both CN and Crz1.

Our data demonstrate that GSH can rescue CN-dependent fitness in the absence of Crz1 (Figure 6). Therefore, we wanted to determine whether GSH rescues fitness due to effects on proliferation, viability, or both. Similar to what we observed in response to CaCl_2_ alone, there was an initial decrease in proliferation rate in all genotypes when cells were grown in monoculture in the presence of both CaCl_2_ and GSH (Figure 7, G-J). In the *CRZ1* background, there was a larger decrease in proliferation rate in *cnb11* cells, similar to what was observed when cells were grown with CaCl_2_ but not GSH. Consistent with the fact that GSH did not rescue fitness in cells with Crz1, the death rates in wild-type and *cnb11* cells were largely unchanged by the addition of GSH (compare Figure 7, A and B with Figure 7, G and H). In contrast, unlike what we observe in response to CaCl_2_ alone (Figure 7, C and D), the proliferation rate decreased to similar extents in both *crz1Δ* and *crz11 cnb11* cells in the presence of CaCl_2_ and GSH and remained constant throughout the time course (Figure 7, I and J). This suggests that GSH rescues the proliferative defect in the *crz11 cnb11* mutant and is consistent with the results of the competitive growth assay (Figure 6A). Additionally, the high CaCl_2_-induced death rate in the *crz11 cnb11* double mutant was decreased in the presence of GSH suggesting that GSH rescues viability in addition to promoting cell cycle progression (Figure 7J).

Finally, we simulated co-culture competitive growth assays in the presence of CaCl_2_ and GSH using proliferation and death rates determined from cells grown in monoculture (Figure 7, K and L). In doing this, we accurately captured the experimentally observed fitness phenotypes of wild-type and *cnb11* cells and confirmed that fitness differences in these strains are driven by proliferation (Figure 7K and Supplemental Figure S6, E and F). Importantly, these simulations also confirmed that GSH has no effect on fitness in the presence of Crz1. In contrast, co-culture simulations of *crz1Δ* and *crz11 cnb11* cells determined that GSH rescues the fitness defect of the double mutant due to effects on proliferation with modest contributions from the GSH-dependent rescue of cell death (Figure 7L and Supplemental Figure S6, G and H). Together, these data demonstrate that reduced proliferation drives the fitness defect in the absence of CN and increased cell death in cells lacking both CN and Crz1 further amplifies this fitness defect.

## DISCUSSION

Coordination between proliferation and stress-defense is critical for long-term survival in challenging environments. The phosphatase CN delays cell cycle progression and promotes gene expression changes to aid adaptation immediately following exposure to CaCl_2_ stress (Yoshimoto *et al*., 2002). However, it was previously unknown whether CN was required for long-term survival in this condition. Here, we set out to examine the impact of CN signaling on fitness during chronic growth in CaCl_2_. This investigation revealed that CN promotes cellular fitness by Crz1-dependent and Crz1-independent mechanisms.

The CN-dependent fitness defect that we describe here has not been previously identified, most likely due to the low sensitivity of standard growth assays. In agreement with previous studies, we did not detect a growth defect in the absence of *CNB1* when cells were grown on agar plates containing CaCl_2_ (Figure 1A) (Nakamura *et al*., 1996; Withee *et al*., 1998; Mulet *et al*., 2006; Ferreira *et al*., 2012; Xu *et al*., 2019). However, cells lacking Cnb1 displayed a proliferative defect in response to CaCl_2_ when they were grown in co-culture with a wild-type strain (Figure 1C). Notably, we did not observe any Crz1-dependent growth defects on agar plates (Figure 1A) despite previous studies reporting sensitivity or lethality in *crz1Δ* cells on agar plates containing CaCl_2_ (Polizotto and Cyert, 2001; Mulet *et al*., 2006; Zhao *et al*., 2013; Xu *et al*., 2019). Though, the loss of Crz1 did result in a competitive fitness defect in the presence of CaCl_2_ (Supplemental Figure S1E) consistent with a previous study (Hsu *et al*., 2021).

Our data demonstrate that CN maintains cellular fitness in CaCl_2_ stress through both Crz1-dependent and Crz1-independent mechanisms (Figure 1). Crz1 is the best characterized downstream target of CN, and many CN-dependent functions and phenotypes are credited to the regulation of this transcription factor (Cyert, 2003; Cyert and Philpott, 2013). However, in addition to Crz1, CN directly regulates many other substrates involved diverse cellular processes including protein trafficking, membrane structure and function, cell cycle, transcription, and translation (Goldman *et al*., 2014). Thus, it is unsurprising that CN impacts fitness in the absence of Crz1.

Although CN can maintain fitness in the absence of Crz1, our data suggests that Crz1 contributes to proper adaptation. CN remains active in cells lacking Crz1 consistent with previous studies demonstrating that Crz1 induces the expression of negative regulators of CN (Figure 2) (Yoshimoto *et al*., 2002). In contrast, when Crz1 is present, most cells inactivate CN within hours of exposure to CaCl_2_ stress suggesting that CN activity is no longer needed to survive, and cells have adapted (Figure 2). Consistent with Crz1 promoting more rapid adaptation, we observed a Crz1-dependent increase in the expression of ribosome biogenesis genes as early as 12 hours after CaCl_2_ treatment (Figures 3 and 4A). As part of the ESR, ribosome biogenesis genes are downregulated in response to stress so that transcription of stress-defense genes can increase (Gasch *et al*., 2000; Causton *et al*., 2001; Levy *et al*., 2007). As adaptation occurs, these gene expression changes are reversed and translational capacity increases so that cells can resume normal growth (Levy *et al*., 2007). We show that both wild type and *cnb11* cells, which express Crz1, increase the expression of ribosome biogenesis genes to enhance translational capacity and this may help cells adapt to chronic CaCl_2_ (Figure 4, A and B). Surprisingly, although there are Crz1-dependent gene expression changes in CN mutants, we see no evidence that our CN reporter, CNR-C, is dephosphorylated or in the nucleus (Figure 2). Since CNR-C is a fragment of Crz1, it is unclear if Crz1 is active in these cells. It is possible that there is a low level of Crz1 activation in a fraction of *cnb11* cells that the reporter is not sensitive enough to detect.

In the absence of Crz1, our data suggest that cells experience redox imbalance and ROS-induced cell death. To prevent ROS accumulation and maintain redox homeostasis cells rely on oxidant defense systems, such as GSH, which act as free radical scavengers or ROS targets (Jamieson, 1998). We found that prolonged CN activity in the absence of Crz1 promotes the upregulation of GSH biosynthesis genes, which helps cells maintain fitness and viability (Figures 5, B and C, and 6). Notably, CN induces the expression of additional genes involved in the oxidative stress response (Yoshimoto *et al*., 2002; Tsuzi *et al*., 2004; Puigpinós *et al*., 2014) and has been shown to suppress cell death in response to other ROS-inducing stressors (Moser *et al*., 1996; Huh *et al*., 2002; Bonilla and Cunningham, 2003; Zhang *et al*., 2006; Dudgeon *et al*., 2008; Farrugia and Balzan, 2012; Zhao *et al*., 2021). However, it is unclear whether CN similarly stimulates GSH production in these conditions. Finally, in mammals, CN promotes autophagy and lysosome biogenesis following activation in response to ROS-induced lysosomal Ca^2+^ release (Zhang *et al*., 2016). These observations suggest that CN may play a conserved role in protecting cells from oxidative damage in response to diverse environmental stressors.

Cells have been shown to secrete GSH in response to environmental stressors such as heat shock and arsenite (Thorsen *et al*., 2012; Trip and Youk, 2020). This raised the possibility that the *crz1Δ* mutant secretes GSH, in addition to upregulating GSH biosynthetic gene expression (Figure 5), which may impact the fitness of *crz11 cnb11* cells when these strains are grown in co-culture. To address this, we determined the proliferation rates and death rates of *crz1Δ* and *crz11 cnb11* mutants growing in monoculture in the presence of CaCl_2_ and used these rates to simulate co-culture experiments (Figure 7, C, D, and F). In doing this, we were able to accurately replicate the experimentally observed co-culture fitness phenotypes (compare Figure 7F and Figure 1E). These findings strongly suggest that if CaCl_2_ does promote GSH secretion, extracellular GSH has negligible, in any, effects on the co-culture fitness of *crz1Δ* and *crz11 cnb11* cells.

Although the immediate response to many environmental stressors, including CaCl_2_, has been well characterized, very little is known about the physiological changes that cells undergo to survive long-term stress exposure. Here, we have characterized the response to chronic CaCl_2_ stress and uncovered CN-dependent mechanisms that allow cells to adapt to this environment. Our findings suggest that the requirements for adaptation change dramatically during prolonged stress exposure compared to the immediate response to stress. This is exemplified in the transcriptional differences we observe in ESR genes either immediately following CaCl_2_ treatment or after many hours in this condition (Supplemental Figure S4D). Additionally, our data indicate that long-term adaptation is stress-specific since prolonged treatment with CaCl_2_ and KCl elicit distinct transcriptional responses (Supplemental Figure S4, A-C). Future studies investigating long-term adaptation to diverse conditions is needed to define the common features of the chronic environmental stress response.

## MATERIALS AND METHODS

### Yeast Strains and Growth Conditions

A complete list of strains used in this study is included in Supplemental Table S1. All strains containing deletions and/or epitope tags were generated using standard methods, as previously described (Rothstein, 1991; Longtine *et al*., 1998). Strains were grown at 30°C in synthetic complete media (C media with 2% dextrose, 1% ammonium chloride). Where indicated, media also contained 0.2 M CaCl_2_, 0.4 M KCl, and/or 250 μM reduced L-glutathione (Sigma), 250 μM DL-homocysteine (Sigma), or 5 mM ascorbic acid (Sigma). Marker strains for competitive fitness assays (GFP+ and GFP-) were constructed as previously described (Conti *et al*., 2022). Any additional genetic manipulations in competition strains (i.e., gene deletions) were introduced by genetic cross between the marker strains and strains containing the desired mutation. Strains in which multiple genes were deleted with the same drug resistance cassette were confirmed by polymerase chain reaction (PCR). All experiments were performed in biological and technical triplicate with independently derived isolates of each strain.

### Serial Dilution Assays

Cells were grown to logarithmic phase growth in synthetic complete media. Equivalent optical densities (0.4 OD_600_) of each sample were collected, washed, and resuspended in 1 mL of synthetic complete media. Five-fold dilutions of strains with the indicated genotypes were then plated on synthetic complete agar (C agar with 2% dextrose and 1% ammonium chloride) or synthetic complete agar containing 0.2 M CaCl_2_ at the indicated concentrations. Plates were incubated at 30°C and imaged after 48 - 72 hours of growth. For experiments using cycloheximide (CHX), ten-fold serial dilutions were plated on YPD agar (2% dextrose) containing the indicated concentrations of CHX and images after 1-8 days of growth at 30°C.

### Competitive Fitness Assays

Co-cultures were generated by mixing equal proportions (1 OD_600_) of cells expressing fluorescent GFP and non-fluorescent GFP-Y66F in 10 mL of synthetic complete media. Immediately following mixing, approximately 0.15 optical densities were collected, washed, resuspended in 2 mL of sodium citrate buffer (50 mM sodium citrate, 0.02% NaN_3_, pH 7.4), and stored at 4°C for subsequent flow cytometric analysis. All co-cultures were then diluted to 0.004-0.01 optical densities in 10 mL of synthetic complete media with or without stress (0.2 M CaCl_2_ or 0.4 M KCl) and incubated at 30°C with continuous rolling. Where indicated, media was supplemented with 250 μM reduced L-glutathione (Sigma), 250 μM DL-homocysteine (Sigma), or 5 mM L-ascorbic acid (Sigma). Co-cultures were sampled and diluted to 0.004-0.02 optical densities, as described above, every 12 hours to ensure that the population did not exceed logarithmic growth during the duration of the experiment. Following sonication, the percentage of GFP and GFP (Y66F) at each timepoint was quantified using a Guava EasyCyte HT flow cytometer and analyzed with FlowJo software. Replicates in which a fitness difference was observed between the GFP and GFP (Y66F) controls were excluded from the final analysis.

### Analysis of Cell Cycle by Flow Cytometry

Approximately 0.15 ODs of cells were fixed in 70% ethanol at 4°C overnight. Following sonication, cells were treated with 0.25 mg/mL RNase A (Biomatik) at 50°C for 1 hour and 0.125 mg/mL Proteinase K (Biomatik) for an additional hour at 50°C. Following deproteination, samples were stained with 1 μM Sytox Green (Invitrogen). DNA content was measured using a Guava EasyCyte HT flow cytometer (Millipore) and analyzed with FlowJo software (FlowJo, LLC).

### CNR-C Imaging

Activation of calcineurin was monitored using a C-terminally GFP-tagged Crz1 truncation which lacks a DNA binding domain (CNR-C). Nuclear localization following dephosphorylation of CNR-C was used as a proxy for *in vivo* calcineurin activity. Following exposure to 0.2 M CaCl_2_ for the indicated times, equivalent optical densities (1 OD_600_) of cells were collected, centrifuged, and resuspended in a small volume of synthetic complete media. Cells were imaged with equivalent exposure times on a Zeiss AxioObserver 7 microscope with a Hamamatsu Orca Fusion-BT camera. ImageJ was used to determine the percentage of cells in the population with nuclear CNR-C signal from a minimum of 100 cells per sample at each timepoint.

### Western Blots

Equivalent optical densities (1 OD_600nm_) of cells were lysed in cold TCA buffer (10 mM Tris-HCl pH 8.0, 10% trichloroacetic acid, 25 mM NH_4_OAc, 1 mM Na_2_EDTA) and incubated on ice. After centrifugation, pellets were resuspended in resuspension solution (0.1M Tris-HCl pH 11, 0.3% SDS) and boiled for 5 minutes at 95°C. Following additional centrifugation, supernatants from clarified samples were transferred to new tubes containing 4X sample buffer (250mM Tris-HCl pH 6.8, 8% SDS, 40% glycerol, 20% β-mercaptoethanol) and boiled for 5 minutes at 95°C. Western blotting was performed using antibodies against GFP (clone JL8, 63268; Clontech), Y15-phosphorylated Cdk1 (anti-Phospho-cdc2, 9111L, Cell Signaling Technology), and PSTAIRE (P7962; Sigma).

### RNA-seq and data analysis

Total RNA was purified using an acid-phenol extraction as previously described (Schmitt *et al*., 1990) from 5 OD_600_ of cells. Poly-A enrichment, library preparation, and sequencing was performed by BGI Genomic Services. Three biological replicates of each time course were performed. All data is available in NCBI GEO and is accessible through GEO Series accession number GSE254555.

RNA-seq analysis was performed with OneStopRNAseq (Li *et al*., 2020). Paired-end reads were aligned to Saccharomyces_cerevisiae.R64-1-1, with star_2.5.3a (Dobin *et al*., 2013), annotated with Saccharomyces_cerevisiae.R64-1-1.90.gtf. Aligned exon fragments with mapping quality higher than 20 were counted toward gene expression with featureCounts (Liao *et al*., 2014). Differential expression (DE) analysis was then performed with DESeq2 (Love *et al*., 2014). Within DE analysis ‘ashr’ was used to create log2 Fold Change (LFC) shrinkage for each comparison (Stephens, 2017). Significant DE genes (DEGs) were identified with the criteria FDR < 0.05. Gene set enrichment analysis was performed with GSEA (Subramanian *et al*., 2005) on the ranked LFC. Significant DE genes were classified into two groups with R package ‘hclust’ using the complete linkage method, with 1 minus Pearson correlation as the distance. Enrichment analysis on each cluster of genes was processed with PANTHER (Thomas *et al*., 2022).

### Quantitation of Petite Frequency

Cells were grown for the indicated times in media containing 0.2 M CaCl_2_. Serial dilutions were performed after collecting 0.1 – 0.2 optical densities of each culture to 1:1000. Equivalent volumes of each diluted culture were plated on YPD agar (2 % dextrose) or YPG (3% glycerol) such that a few hundred colonies were formed. YPD and YPG plates were imaged after 3 and 7 days of growth at 30°C, respectively. For each genotype, petite frequency was determined from n = 6 biological replicates in which at least 200 colonies grew on YPD agar using the following formula:

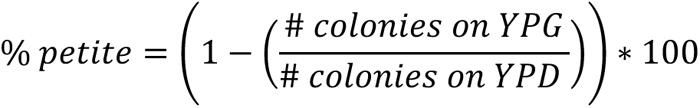

### Propidium Iodide Staining

After sample collection and centrifugation, approximately 0.15 ODs of cells were resuspended in 2 mL sodium citrate buffer (50 mM sodium citrate, 0.02% NaN_3_, pH 7.4) and stored at 4°C. Following washing with additional sodium citrate buffer, propidium iodide (PI) (MP Biomedicals, LLC) was added to each sample to a final concentration of 3 μg/mL. Samples were then incubated in the dark for 30 minutes prior to sonication. The percentage of PI positive cells were quantified using a Guava EasyCyte HT flow cytometer and analyzed with FlowJo software.

### Calculation of Proliferation and Death Rates

For each experimental condition, doubling times of strains grown in monoculture were calculated by fitting OD_600_ values obtained at 12-hour intervals to an exponential growth equation, where *G* = number of generations, *P_0_* = initial population size, and *P_t_* = final population size:

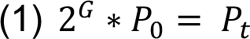

Doubling times were then determined using the following equation:

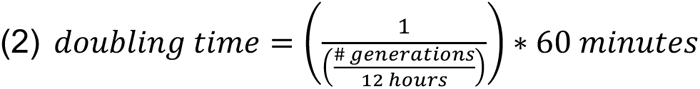

Importantly, the observed doubling time reflects the net effect, combining the true proliferation rate and true death rate. True proliferation rates and death rates were calculated using the GRADE method, as previously described (Schwartz *et al*., 2020). Briefly, the GRADE method infers the true cell proliferation rate from a combination of 1) the experimentally measured death rate, and 2) the experimentally measured net population growth rate. The average death rate was determined using flow cytometry-based measurement of percent propidium iodide positivity (described above), to determine the “fractional viability” (FV) where *C_live_* = live cell number, *C_dead_* = dead cell number:

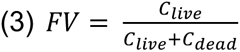

The net population growth rate (GR value) was computed as previously described (Hafner *et al*., 2016), using relative OD_600_ values, where *C_0_*= initial cell number, *C_live_*= live cell number, *C_dead_* = dead cell number, and *C_ctrl_* = live cell number of a reference condition (defined here as a population with maximum proliferation rate and a death rate of zero):

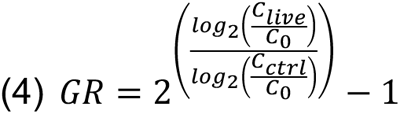

To identify the true proliferation rate and death rate, we simulated all pairwise combinations of 500 proliferation rates and 500 death rates using the following equations, where *C_0_* = initial cell number, *t* = assay duration, τ = proliferation rate, and *D_R_* = death rate.

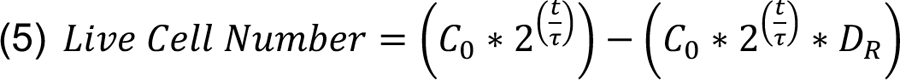

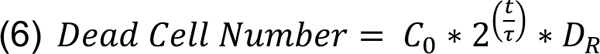

This yielded average population dynamics for 250,000 plausible proliferation rate and death rate pairs. For each simulated proliferation rate and death rate pair, the fractional viability (FV) value and growth rate inhibition (GR) value (Hafner *et al*., 2016)were calculated, as defined above. Each simulated proliferation rate and death rate pair are uniquely identifiable by their FV and GR values. Thus, the true proliferation rates and death rates of the experimental conditions were then determined by finding which simulation best matches the observed FV/GR pair.

*In silico* co-culture experiments (Figures 7, E,F,K, and L, and Supplemental Figure 6) were simulated using the experimentally determined proliferation rates and death rates from monoculture (described above) as follows. Two genotypes were seeded at equal cell numbers at time zero. We assumed there was no interaction between genotypes. Total live cell numbers for each genotype evolved according to equation (5), and live cell numbers, proliferation rates, and death rates were updated at each experimental time interval (every 12 hours for 3 days). To separately evaluate the role of proliferation rate and death rate on *in silico* co-culture dynamics, simulations were re-run, setting either the proliferation rates or death rates to be equal between the two genotypes.

## Abbreviations used

CN: calcineurin
ESR: environmental stress response
MAPK: mitogen-activated protein kinase
ROS: reactive oxygen species
GSH: glutathione
ETC: electron transport chain
CHX: cycloheximide
PI: propidium iodide

## ACKNOWLEDGEMENTS

We thank Tom Fazzio and members of the Benanti lab for helpful discussions and critical reading of the manuscript. This work was supported by National Institutes of Health grants R35GM136280 to J. A. B., and R01GM127559 to M.J.L.

## REFERENCES

Alberts, B, Bray, D, Lewis, J, Raff, M, Roberts, K, and Watson, JD (1994). Molecular Biology of the Cell 3rd Edition, Garland Publishing Inc.

Bonilla, M, and Cunningham, KW (2003). Mitogen-activated Protein Kinase Stimulation of Ca 2+ Signaling Is Required for Survival of Endoplasmic Reticulum Stress in Yeast. Mol Biol Cell 14, 4296–4305.

Bonny, AR, Kochanowski, K, Diether, M, and El-Samad, H (2021). Stress-induced growth rate reduction restricts metabolic resource utilization to modulate osmo-adaptation time. Cell Reports 34, 108854.

Bothammal, P, Prasad, M, Muralitharan, G, and Natarajaseenivasan, K (2022). Leptospiral lipopolysaccharide mediated Hog1 phosphorylation in Saccharomyces cerevisiae directs activation of autophagy. Microb Pathog 173, 105840.

Brennan, RJ, and Schiestl, RH (1996). Cadmium is an inducer of oxidative stress in yeast. Mutat ResFundam Mol Mech Mutagen 356, 171–178.

Carmona-Gutierrez, D, Bauer, MA, Zimmermann, A, Aguilera, A, Austriaco, N, Ayscough, K, Balzan, R, Bar-Nun, S, Barrientos, A, Belenky, P, et al. (2018). Guidelines and recommendations on yeast cell death nomenclature. Microb Cell 5, 4.

Carraro, M, and Bernardi, P (2016). Calcium and reactive oxygen species in regulation of the mitochondrial permeability transition and of programmed cell death in yeast. Cell Calcium 60, 102–107.

Causton, HC, Ren, B, Koh, SS, Harbison, CT, Kanin, E, Jennings, EG, Lee, TI, True, HL, Lander, ES, and Young, RA (2001). Remodeling of Yeast Genome Expression in Response to Environmental Changes. Mol Biol Cell 12, 323–337.

Clotet, J, Escoté, X, Adrover, MÀ, Yaakov, G, Garí, E, Aldea, M, Nadal, E de, and Posas, F (2006). Phosphorylation of Hsl1 by Hog1 leads to a G2 arrest essential for cell survival at high osmolarity. EMBO J 25, 2338–2346.

Conti, MM, Ghizzoni, JM, Gil-Bona, A, Wang, W, Costanzo, M, Li, R, Flynn, MJ, Zhu, LJ, Myers, CL, Boone, C, et al. (2022). Repression of essential cell cycle genes increases cellular fitness. PLoS Genet 18, e1010349.

Cyert, MS (2003). Calcineurin signaling in Saccharomyces cerevisiae: how yeast go crazy in response to stress. Biochem Biophys Res Commun 311, 1143–1150.

Cyert, MS, and Philpott, CC (2013). Regulation of cation balance in Saccharomyces cerevisiae. Genetics 193, 677–713.

Davidson, JF, Whyte, B, Bissinger, PH, and Schiestl, RH (1996). Oxidative stress is involved in heat-induced cell death in Saccharomyces cerevisiae. Proc Natl Acad Sci 93, 5116–5121.

Deere, D, Shen, J, Vesey, G, Bell, P, Bissinger, P, and Veal, D (1998). Flow cytometry and cell sorting for yeast viability assessment and cell selection. Yeast 14, 147–160.

Dobin, A, Davis, CA, Schlesinger, F, Drenkow, J, Zaleski, C, Jha, S, Batut, P, Chaisson, M, and Gingeras, TR (2013). STAR: ultrafast universal RNA-seq aligner. Bioinformatics 29, 15–21.

Drakulic, T, Temple, MD, Guido, R, Jarolim, S, Breitenbach, M, Attfield, PV, and Dawes, IW (2005). Involvement of oxidative stress response genes in redox homeostasis, the level of reactive oxygen species, and ageing in Saccharomyces cerevisiae. FEMS Yeast Res 5, 1215– 1228.

Du, L, Yu, Y, Chen, J, Liu, Y, Xia, Y, Chen, Q, and Liu, X (2007). Arsenic induces caspase- and mitochondria-mediated apoptosis in Saccharomyces cerevisiae. FEMS Yeast Res 7, 860–865.

Dudgeon, DD, Zhang, N, Ositelu, OO, Kim, H, and Cunningham, KW (2008). Nonapoptotic Death of Saccharomyces cerevisiae Cells That Is Stimulated by Hsp90 and Inhibited by Calcineurin and Cmk2 in Response to Endoplasmic Reticulum Stresses. Eukaryot Cell 7, 2037– 2051.

Elliott, B, and Futcher, B (1993). Stress resistance of yeast cells is largely independent of cell cycle phase. Yeast 9, 33–42.

Elskens, MT, Jaspers, CJ, and Penninckx, MJ (1991). Glutathione as an endogenous sulphur source in the yeast Saccharomyces cerevisiae. Microbiology 137, 637–644.

Epstein, CB, Waddle, JA, Hale, W, Davé, V, Thornton, J, Macatee, TL, Garner, HR, and Butow, RA (2001). Genome-wide Responses to Mitochondrial Dysfunction. Mol Biol Cell 12, 297–308.

Farrugia, G, and Balzan, R (2012). Oxidative Stress and Programmed Cell Death in Yeast. Front Oncol 2, 64.

Ferreira, RT, Silva, ARC, Pimentel, C, Batista-Nascimento, L, Rodrigues-Pousada, C, and Menezes, RA (2012). Arsenic stress elicits cytosolic Ca2+ bursts and Crz1 activation in Saccharomyces cerevisiae. Microbiology 158, 2293–2302.

Flynn, MJ, and Benanti, JA (2022). Cip1 tunes cell cycle arrest duration upon calcineurin activation. Proc Natl Acad Sci United States Am 119, e2202469119.

Frigo, E, Tommasin, L, Lippe, G, Carraro, M, and Bernardi, P (2023). The Haves and Have-Nots: The Mitochondrial Permeability Transition Pore across Species. Cells 12, 1409.

Gasch, AP, Spellman, PT, Kao, CM, Carmel-Harel, O, Eisen, MB, Storz, G, Botstein, D, and Brown, PO (2000). Genomic Expression Programs in the Response of Yeast Cells to Environmental Changes. Mol Biol Cell 11, 4241–4257.

Goldman, A, Roy, J, Bodenmiller, B, Wanka, S, Landry, CR, Aebersold, R, and Cyert, MS (2014). The Calcineurin Signaling Network Evolves via Conserved Kinase-Phosphatase Modules that Transcend Substrate Identity. Mol Cell 55, 422–435.

Granot, D, Levine, A, and Dor-Hefetz, E (2003). Sugar-induced apoptosis in yeast cells. FEMS Yeast Res 4, 7–13.

Guaragnella, N, Ždralević, M, Antonacci, L, Passarella, S, Marra, E, and Giannattasio, S (2012). The role of mitochondria in yeast programmed cell death. Front Oncol 2, 70.

Hafner, M, Niepel, M, Chung, M, and Sorger, PK (2016). Growth rate inhibition metrics correct for confounders in measuring sensitivity to cancer drugs. Nat Methods 13, 521–527.

Ho, Y-H, and Gasch, AP (2015). Exploiting the yeast stress-activated signaling network to inform on stress biology and disease signaling. Current Genetics 61, 503–511.

Hsu, IS, Strome, B, Lash, E, Robbins, N, Cowen, LE, and Moses, AM (2021). A functionally divergent intrinsically disordered region underlying the conservation of stochastic signaling. Plos Genet 17, e1009629.

Huh, G, Damsz, B, Matsumoto, TK, Reddy, MP, Rus, AM, Ibeas, JI, Narasimhan, ML, Bressan, RA, and Hasegawa, PM (2002). Salt causes ion disequilibrium-induced programmed cell death in yeast and plants. Plant J 29, 649–659.

Jamieson, DJ (1998). Oxidative stress responses of the yeast Saccharomyces cerevisiae. Yeast 14, 1511–1527.

Leech, CM, Flynn, MJ, Arsenault, HE, Ou, J, Liu, H, Zhu, LJ, and Benanti, JA (2020). The coordinate actions of calcineurin and Hog1 mediate the stress response through multiple nodes of the cell cycle network. PLoS Genetics 16, e1008600–27.

Levy, S, Ihmels, J, Carmi, M, Weinberger, A, Friedlander, G, and Barkai, N (2007). Strategy of Transcription Regulation in the Budding Yeast. PLoS ONE 2, e250.

Li, R, Hu, K, Liu, H, Green, MR, and Zhu, LJ (2020). OneStopRNAseq: A Web Application for Comprehensive and Efficient Analyses of RNA-Seq Data. Genes 11, 1165.

Liao, Y, Smyth, GK, and Shi, W (2014). featureCounts: an efficient general purpose program for assigning sequence reads to genomic features. Bioinformatics 30, 923–930.

Longtine, MS, McKenzie, A, Demarini, DJ, Shah, NG, Wach, A, Brachat, A, Philippsen, P, and Pringle, JR (1998). Additional modules for versatile and economical PCR-based gene deletion and modification in Saccharomyces cerevisiae. Yeast 14, 953–961.

López-Maury, L, Marguerat, S, and Bähler, J (2008). Tuning gene expression to changing environments: from rapid responses to evolutionary adaptation. Nat Rev Genet 9, 583–593.

Love, MI, Huber, W, and Anders, S (2014). Moderated estimation of fold change and dispersion for RNA-seq data with DESeq2. Genome Biol 15, 550.

Lu, C, Brauer, MJ, and Botstein, D (2008). Slow growth induces heat-shock resistance in normal and respiratory-deficient yeast. Mol Biol Cell 20, 891–903.

Matheos, DP, Kingsbury, TJ, Ahsan, US, and Cunningham, KW (1997). Tcn1p/Crz1p, a calcineurin-dependent transcription factor that differentially regulates gene expression in Saccharomyces cerevisiae. Genes Dev 11, 3445–3458.

Medina, DL, Paola, SD, Peluso, I, Armani, A, Stefani, DD, Venditti, R, Montefusco, S, Scotto-Rosato, A, Prezioso, C, Forrester, A, et al. (2015). Lysosomal calcium signalling regulates autophagy through calcineurin and TFEB. Nat Cell Biol 17, 288–299.

Mirisola, MG, Braun, RJ, and Petranovic, D (2014). Approaches to study yeast cell aging and death. FEMS Yeast Res 14, 109–118.

Mizunuma, M, Hirata, D, Miyahara, K, Tsuchiya, E, and Miyakawa, T (1998). Role of calcineurin and Mpk1 in regulating the onset of mitosis in budding yeast. Nature 392, 303–306.

Mizunuma, M, Hirata, D, Miyaoka, R, and Miyakawa, T (2001). GSK-3 kinase Mck1 and calcineurin coordinately mediate Hsl1 down-regulation by Ca2+ in budding yeast. EMBO J 20, 1074–1085.

Moser, MJ, Geiser, JR, and Davis, TN (1996). Ca2+-Calmodulin Promotes Survival of Pheromone-Induced Growth Arrest by Activation of Calcineurin and Ca2+-Calmodulin-Dependent Protein Kinase. Mol Cell Biol 16, 4824–4831.

Mulet, JM, Martin, DE, Loewith, R, and Hall, MN (2006). Mutual Antagonism of Target of Rapamycin and Calcineurin Signaling*. J Biol Chem 281, 33000–33007.

Nakamura, T, Ohmoto, T, Hirata, D, Tsuchiya, E, and Miyakawa, T (1996). Genetic evidence for the functional redundancy of the calcineurin-and Mpk1-mediated pathways in the regulation of cellular events important for growth inSaccharomyces cerevisiae. Mol Gen Genet MGG 251, 211–219.

Polizotto, RS, and Cyert, MS (2001). Calcineurin-dependent nuclear import of the transcription factor Crz1p requires Nmd5p. J Cell Biol 154, 951–960.

Puigpinós, J, Casas, C, and Herrero, E (2014). Altered intracellular calcium homeostasis and endoplasmic reticulum redox state in Saccharomyces cerevisiae cells lacking Grx6 glutaredoxin. Mol Biol Cell 26, mbc.E14-06–1137.

Rothstein, R (1991). Targeting, disruption, replacement, and allele rescue: integrative DNA transformation in yeast. Methods in Enzymology 194, 281–301.

Schmitt, ME, Brown, TA, and Trumpower, BL (1990). A rapid and simple method for preparation of RNA from Saccharomyces cerevisiae. Nucleic Acids Res 18, 3091–3092.

Schwartz, HR, Richards, R, Fontana, RE, Joyce, AJ, Honeywell, ME, and Lee, MJ (2020). Drug GRADE: An Integrated Analysis of Population Growth and Cell Death Reveals Drug-Specific and Cancer Subtype-Specific Response Profiles. Cell Rep 31, 107800.

Stathopoulos, AM, and Cyert, MS (1997). Calcineurin acts through theCRZ1/TCN1-encoded transcription factor to regulate gene expression in yeast. Genes Dev 11, 3432–3444.

Stathopoulos-Gerontides, A, Guo, JJ, and Cyert, MS (1999). Yeast calcineurin regulates nuclear localization of the Crz1p transcription factor through dephosphorylation. Genes Dev 13, 798– 803.

Stephens, M (2017). False discovery rates: a new deal. Biostat Oxf Engl 18, 275–294.

Subramanian, A, Tamayo, P, Mootha, VK, Mukherjee, S, Ebert, BL, Gillette, MA, Paulovich, A, Pomeroy, SL, Golub, TR, Lander, ES, et al. (2005). Gene set enrichment analysis: A knowledge-based approach for interpreting genome-wide expression profiles. Proc Natl Acad Sci 102, 15545–15550.

Thomas, PD, Ebert, D, Muruganujan, A, Mushayahama, T, Albou, L, and Mi, H (2022). PANTHER: Making genome-scale phylogenetics accessible to all. Protein Sci 31, 8–22.

Thorsen, M, Jacobson, T, Vooijs, R, Navarrete, C, Bliek, T, Schat, H, and Tamás, MJ (2012). Glutathione serves an extracellular defence function to decrease arsenite accumulation and toxicity in yeast. Mol Microbiol 84, 1177–1188.

Trip, DSL, Maire, T, and Youk, H (2022). Fundamental limits to progression of cellular life in frigid environments. BioRxiv, 2022.06.10.495632.

Trip, DSL, and Youk, H (2020). Yeasts collectively extend the limits of habitable temperatures by secreting glutathione. Nat Microbiol 5, 943–954.

Tsuzi, D, Maeta, K, Takatsume, Y, Izawa, S, and Inoue, Y (2004). Distinct regulatory mechanism of yeast GPX2 encoding phospholipid hydroperoxide glutathione peroxidase by oxidative stress and a calcineurin/Crz1-mediated Ca2+ signaling pathway. FEBS Lett 569, 301– 306.

Withee, JL, Mulholland, J, Jeng, R, and Cyert, MS (1997). An essential role of the yeast pheromone-induced Ca2+ signal is to activate calcineurin. Mol Biol Cell 8, 263–277.

Withee, JL, Sen, R, and Cyert, MS (1998). Ion Tolerance of Saccharomyces cerevisiae Lacking the Ca2+/CaM-Dependent Phosphatase (Calcineurin) Is Improved by Mutations in URE2 or PMA1. Genetics 149, 865–878.

Xu, H, Fang, T, Yan, H, and Jiang, L (2019). The protein kinase Cmk2 negatively regulates the calcium/calcineurin signalling pathway and expression of calcium pump genes PMR1 and PMC1 in budding yeast. Cell Commun Signal 17, 7.

Yi, D-G, Hong, S, and Huh, W-K (2018). Mitochondrial dysfunction reduces yeast replicative lifespan by elevating RAS-dependent ROS production by the ER-localized NADPH oxidase Yno1. PLoS ONE 13, e0198619.

Yokoyama, H, Mizunuma, M, Okamoto, M, Yamamoto, J, Hirata, D, and Miyakawa, T (2006). Involvement of calcineurin-dependent degradation of Yap1p in Ca2+-induced G2 cell-cycle regulation in Saccharomyces cerevisiae. EMBO Rep 7, 519–524.

Yoshimoto, H, Saltsman, K, Gasch, AP, Li, HX, Ogawa, N, Botstein, D, Brown, PO, and Cyert, MS (2002). Genome-wide Analysis of Gene Expression Regulated by the Calcineurin/Crz1p Signaling Pathway in Saccharomyces cerevisiae *. J Biol Chem 277, 31079–31088.

Zakrzewska, A, Eikenhorst, G van, Burggraaff, JEC, Vis, DJ, Hoefsloot, H, Delneri, D, Oliver, SG, Brul, S, and Smits, GJ (2011). Genome-wide analysis of yeast stress survival and tolerance acquisition to analyze the central trade-off between growth rate and cellular robustness. Mol Biol Cell 22, 4435–4446.

Zhang, N-N, Dudgeon, DD, Paliwal, S, Levchenko, A, Grote, E, and Cunningham, KW (2006). Multiple Signaling Pathways Regulate Yeast Cell Death during the Response to Mating Pheromones. Mol Biol Cell 17, 3409–3422.

Zhang, X, Cheng, X, Yu, L, Yang, J, Calvo, R, Patnaik, S, Hu, X, Gao, Q, Yang, M, Lawas, M, et al. (2016). MCOLN1 is a ROS sensor in lysosomes that regulates autophagy. Nat Commun 7, 12109.

Zhao, Y, Du, J, Zhao, G, and Jiang, L (2013). Activation of calcineurin is mainly responsible for the calcium sensitivity of gene deletion mutations in the genome of budding yeast. Genomics 101, 49–56.

Zhao, Y, Su, R, Li, S, and Mao, Y (2021). Mechanistic analysis of cadmium toxicity in Saccharomyces cerevisiae. FEMS Microbiol Lett 368.

