## Supplemental Figures for "Calcineurin promotes adaptation to chronic stress through two distinct mechanisms"

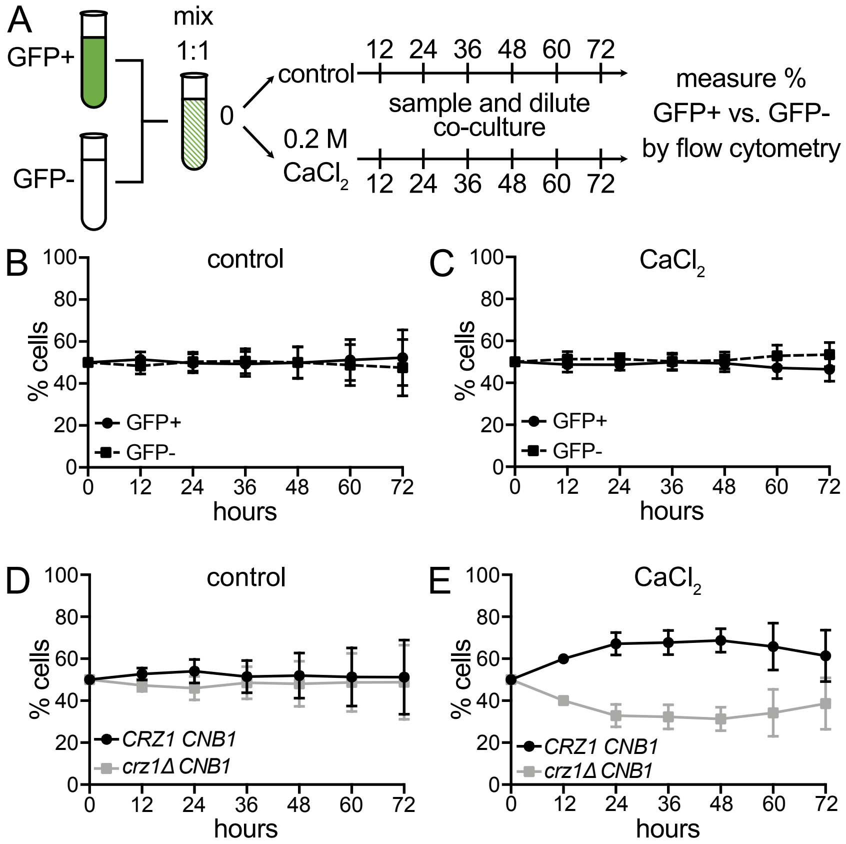


Supplemental Figure S1. Overview of the co-culture competitive fitness assays. (A) Experimental schematic of the co-culture competition assay. A strain expressing wild-type GFP is co-cultured in a 1:1 ratio with a strain expressing a non-fluorescent GFP mutant (Y66F). Co-cultures are sampled and diluted every 12 hours to prevent cells from exiting logarithmic phase growth. The relative proportions of each strain are measured by flow cytometry. (B-C) GFP+ and GFP- strains were co-cultured as in (A) in the absence (B) or presence (C) of 0.2 M $CaCl_2$ and diluted as described in (A). An average of n = 11 biological replicates is shown. Error bars represent standard deviations. (D-E) *CRZ1 CNB1* and *crz1Δ CNB1* cells were co-cultured as described in (A) in the absence (D) or presence (E) of 0.2 M $CaCl_2$. Shown is an average of n = 9 biological replicates. Error bars indicate standard deviations.

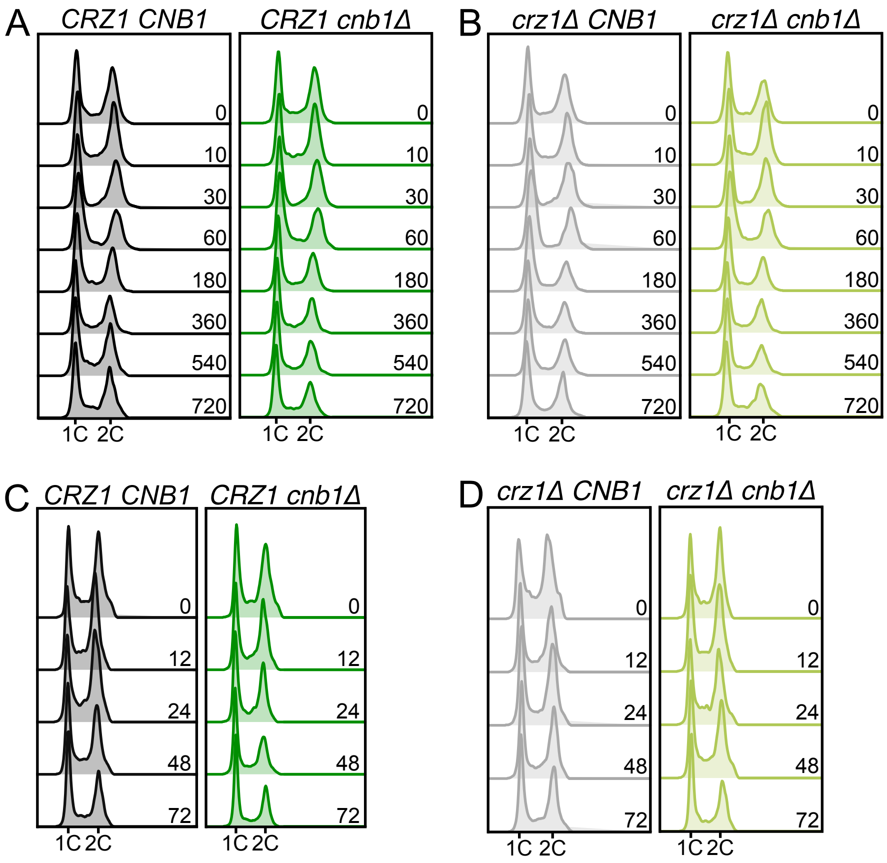


Supplemental Figure S2. Cell cycle progression is altered in the absence of CN. (A) *CRZ1 CNB1* and *CRZ1 cnb1Δ* cells were grown in 0.2 M $CaCl_2$, and DNA content was measured at the indicated times by flow cytometry. Representative histograms from n = 3 experiments are shown. (B) Same as (A) in *crz1Δ CNB1* and *crz1Δ cnb1Δ* cells. Representative histograms from n = 3 experiments are shown. In (A) and (B), a CN-independent decrease in S phase is evident at the 30-minute timepoint and CN-dependent maintence of this S phase decrease is evident until the 60-minute timepoint, as previously described (Leech *et al.*, 2020). (C) *CRZ1 CNB1* and *CRZ1 cnb1Δ* cells were grown in 0.2 M $CaCl_2$ and diluted every 12 hours. Samples were collected at the indicated times and DNA content was measured by flow cytometry. Representative histograms from n = 3 experiments are shown. (D) Same as (C) in *crz1Δ CNB1* and *crz1Δ cnb1Δ* cells. Representative histograms from n = 3 experiments are shown. In (C) and (D) modest CN-dependent differences in overall cell cycle profiles are evident after 12 to 72 hours of $CaCl_2$ treatment.

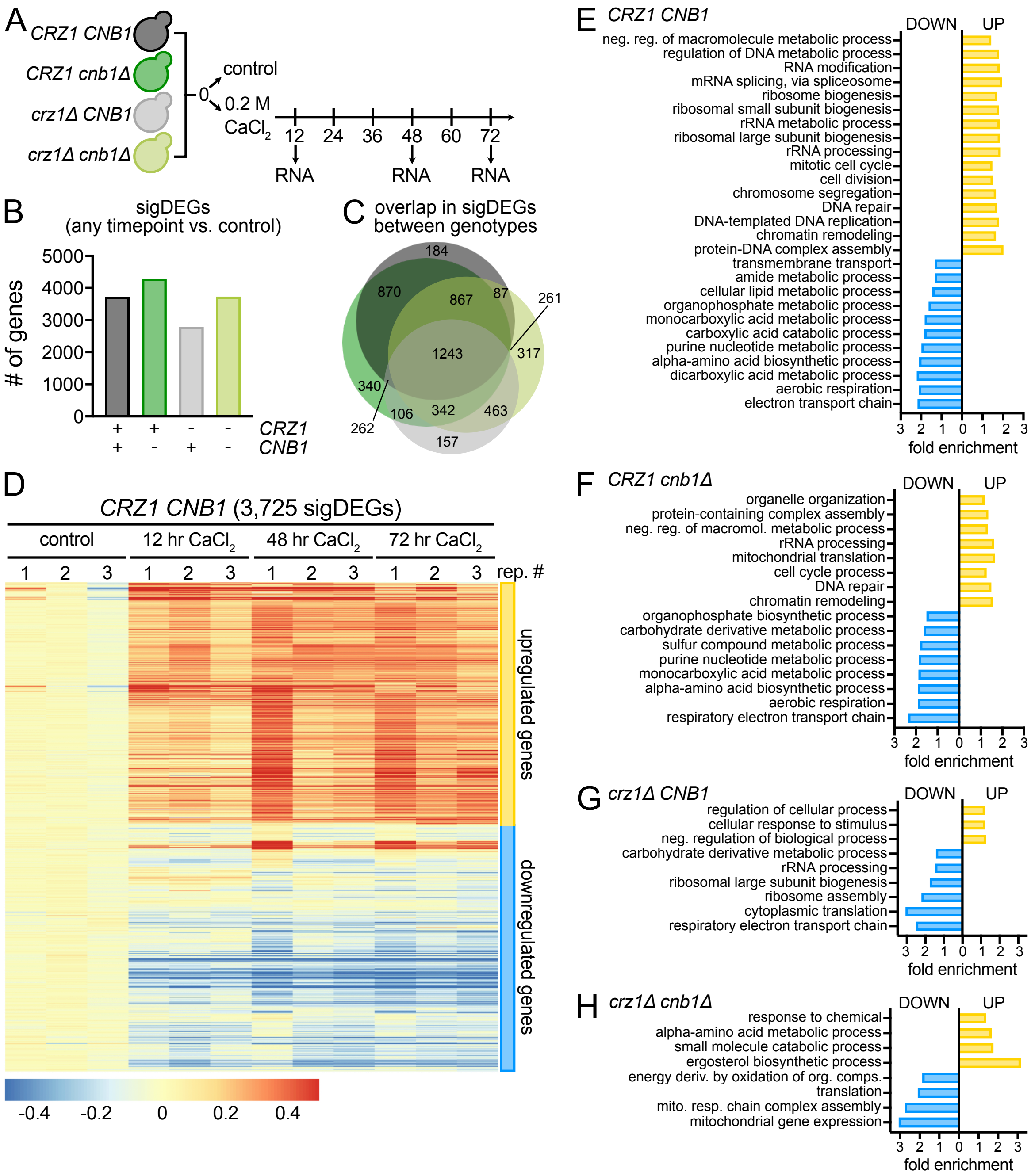


Supplemental Figure S3. Overview of RNA-seq experiment and GO term summary. (A) Experimental schematic of RNA-seq experiment. Cells were grown in 0.2 M $CaCl_2$ for the indicated times and RNA was isolated after 12, 48, and 72 hours in $CaCl_2$ for RNA-seq. RNA from cells grown in the absence of $CaCl_2$ was used as a control for the differential gene expression analysis. (B) Summary of DESeq2 results. The number of genes that were significantly differentially expressed (sigDEGs) (FDR < 0.05, fold change > 0.585) at any timepoint compared to an unstressed control for each genotype is shown. (C) Overlap of all sigDEGs across the indicated genotypes. (D) Hierarchical clustering of sigDEGs used for Gene Ontology (GO)-term analysis in *CRZ1 CNB1* cells. The expression value heat map shows the $\log_{10}$(TPM+1) centered around the mean of the control group, from n = 3 biological replicates. Up- and downregulated clusters are indicated on the right. Heatmaps for additional genotypes can be found in Supplemental Data File S2. (E) A selection of significantly enriched GO-terms in *CRZ1 CNB1* cells. Yellow and blue bars indicate processes that were enriched in upregulated or downregulated genes from (D), respectively. (F-H) Same as (E) in *CRZ1 cnb1Δ* (F), *crz1Δ CNB1* (G), and *crz1Δ cnb1Δ* (H) cells. See also Supplemental Data File S2.

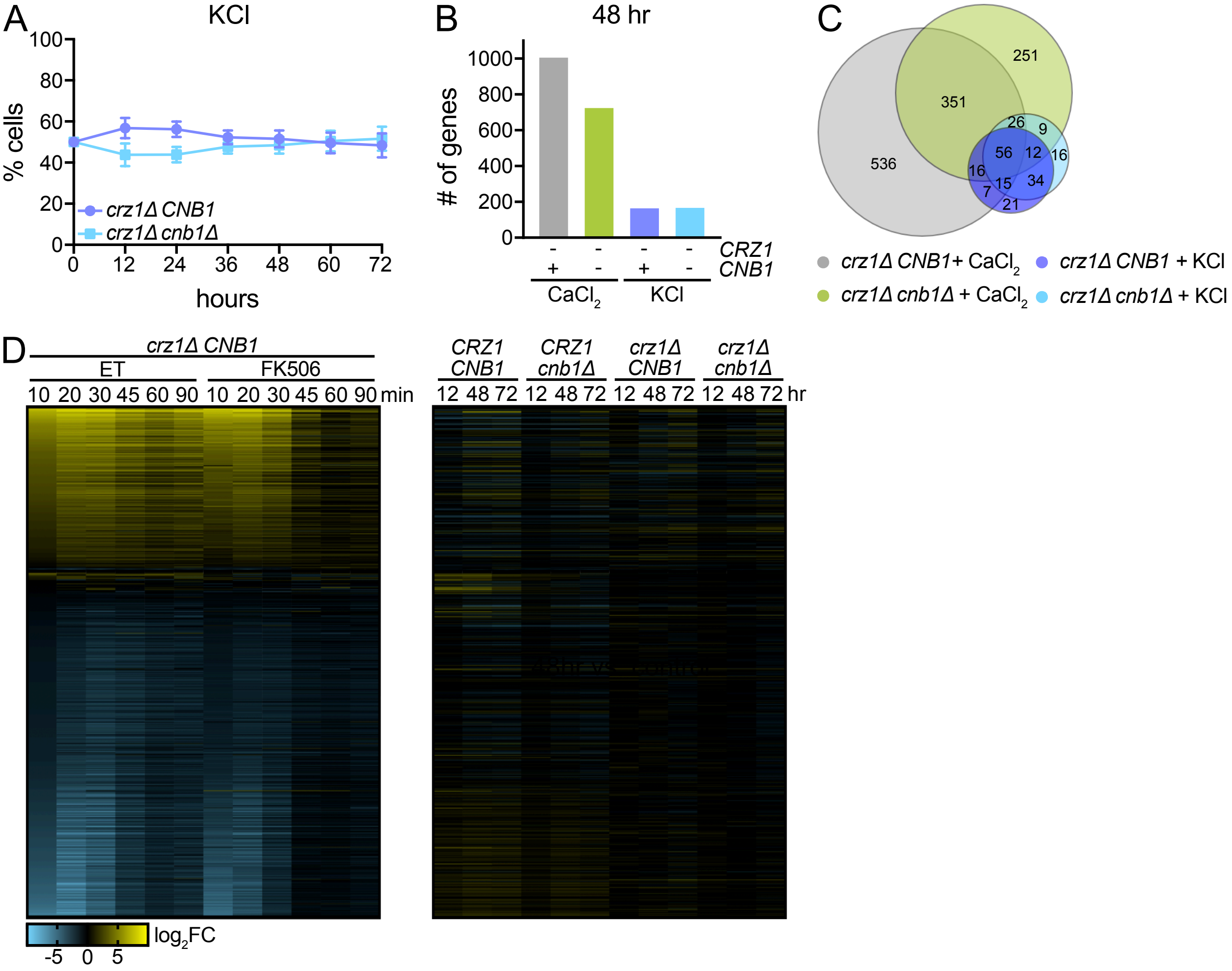


Supplemental Figure S4. The response to chronic KCl and acute $CaCl_2$ stress. (A) *crz1Δ CNB1* and *crz1Δ cnb1Δ* cells were co-cultured in the presence of 0.4 M KCl. Co-cultures were sampled and diluted every 12 hours. The relative proportion of each strain was determined at the indicated timepoints by flow cytometry. An average of n = 9 biological replicates is shown, error bars indicate standard deviations. (B-C) Comparison of the number (B) and overlap (C) of significantly differentially expressed genes after 48 hours of growth in 0.2 M $CaCl_2$ or 0.4 M KCl. Lists of genes included can be found in Supplemental Data File S1. (D) Heatmap showing the $\log_2$ fold change in expression of ESR genes (Gasch *et al.,* 2000) in response to 0.2 M acute (left) or chronic (right) $CaCl_2$ in the indicated genotypes. During acute stress, *crz1Δ CNB1* cells were pretreated with either control buffer (ET) or the CN inhibitor (FK506) for 15 minutes before the addition of 0.2 M $CaCl_2$ (Leech *et al.*, 2020). A list of genes and $\log_2$ fold change values used to generate the heatmap are included in Supplemental Data File S4.

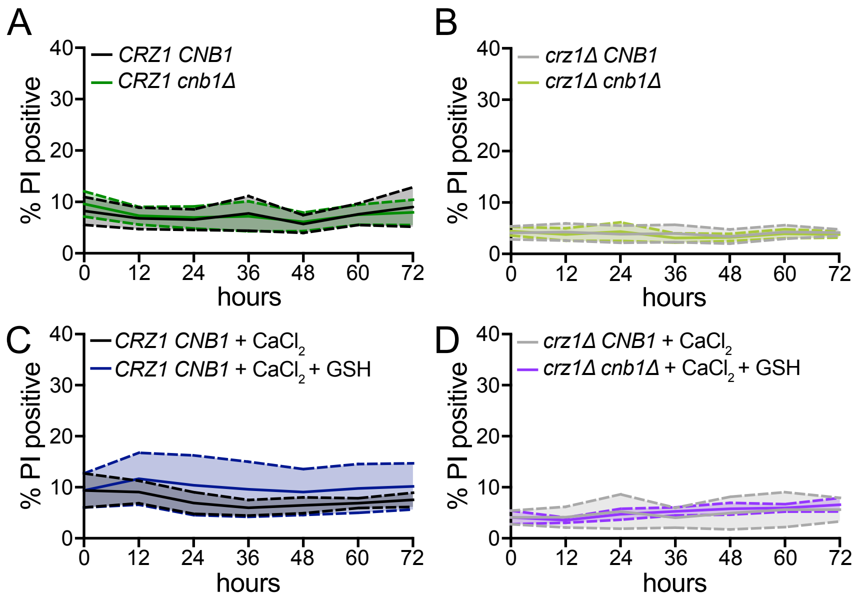


Supplemental Figure S5. Data in support of cell death measurements. (A-B) *CRZ1 CNB1* and *CRZ1 cnb1Δ* (A) and *crz1Δ CNB1* and *crz1Δ cnb1Δ* (B) cells and were grown in monoculture sampled and diluted every 12 hours and the percentage of propidium iodide (PI) positive cells was measured by flow cytometry. An average of n = 12 (A) or n = 9 (B) biological replicates is shown. Solid line denotes the mean and the shaded area between the dashed lines denotes the standard error of the mean. (C-D) As in (A-B) in *CRZ1 CNB1* (C) and *crz1Δ CNB1* (D) cells in the presence of 0.2 M $CaCl_2$ or 0.2 M $CaCl_2$ and 250 µM GSH. An average of n = 9 biological replicates is shown. Solid line denotes the mean and the shaded area within the dashed lines denotes the standard error of the mean.

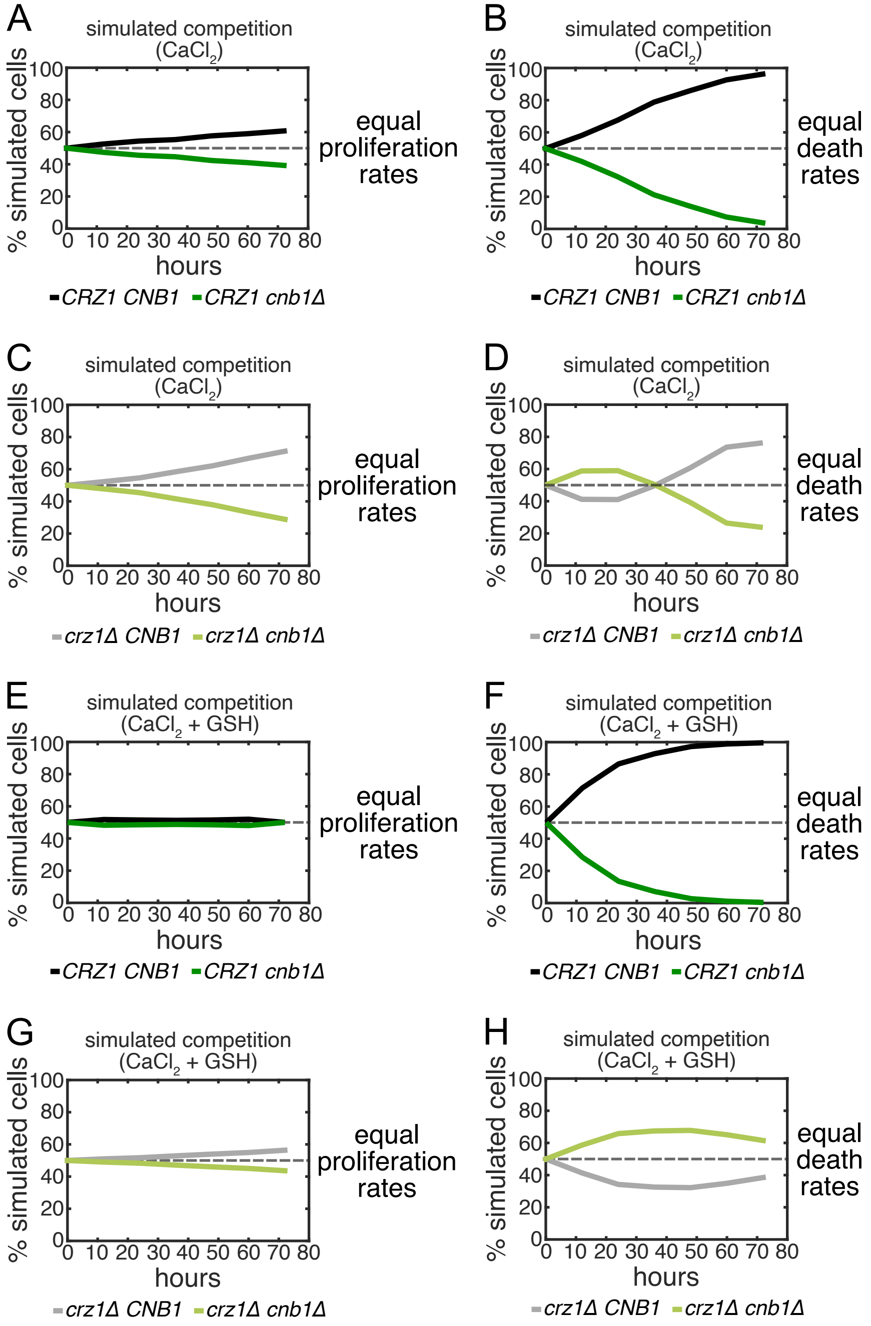


Supplemental Figure S6. Simulated competition assays with fixed proliferation or death rates. (A) Simulated co-culture competition assay between *CRZ1 CNB1* and *CRZ1 cnb1Δ* cells with equal proliferation rates in the presence of 0.2 M $CaCl_2$. (B) Simulated co-culture competition assay between *CRZ1 CNB1* and *CRZ1 cnb1Δ* cells with equal death rates in the presence of 0.2 M $CaCl_2$.(C) Same as (A) in *crz1Δ CNB1* and *crz1Δ cnb1Δ* cells. (D) Same as (B) in *crz1Δ CNB1* and *crz1Δ cnb1Δ* cells. (E) Simulated co-culture competition assay between *CRZ1 CNB1* and *CRZ1 cnb1Δ* cells with equal proliferation rates in the presence of 0.2 M $CaCl_2$ and 250 μM GSH. (F) Simulated co-culture competition assay between *CRZ1 CNB1* and *CRZ1 cnb1Δ* cells with equal death rates in the presence of 0.2 M $CaCl_2$ and 250 μM GSH. (G) Same as (E) in *crz1Δ CNB1* and *crz1Δ cnb1Δ* cells. (H) Same as (F) in *crz1Δ CNB1* and *crz1Δ cnb1Δ* cells.
